# Brush border intermicrovillar adhesion limits bacteria attachment to the small intestine brush border

**DOI:** 10.1101/2025.11.25.690363

**Authors:** Rebecca P. Cowell, Alexander C. Bickart, Suchetha R. Shashi, Alfredo Vasquez, Lee C. Brackman, Marwa A. Abdel-Dayem, Wei Chen, Deanna M. Bowman, Evan S. Krystofiak, Yassine Essalki, Isabelle Gross, David G. Harrison, Celestine N. Wanjalla, Holly M. Scott Algood, Matthew J. Tyska, Leslie M. Meenderink

**Affiliations:** Division of Infectious Disease, Department of Medicine, Vanderbilt University School of Medicine, Vanderbilt University Medical Center, Nashville, TN, USA; Tennessee Valley Healthcare System, Department of Veterans Affairs, Nashville, TN, USA; Division of Genetic Medicine and Clinical Pharmacology, Department of Medicine, Vanderbilt University Medical Center, Nashville, TN, USA; Department of Biochemistry and Pharmacology, Faculty of Pharmacy, Horus University-Egypt, New Damietta 34518, Egypt; College of Graduate Studies, Northeast Ohio Medical School, Rootstown, OH, USA; Department of Cell and Developmental Biology, Vanderbilt University School of Medicine, Nashville, TN, USA; Regenerative Nanomedicine, INSERM UMR 1260, University of Strasbourg, CRBS, FMTS, Strasbourg, France; Tennessee Center for AIDS Research, Vanderbilt University Medical Center, Nashville, TN, USA; Department of Pathology, Microbiology and Immunology, Vanderbilt University School of Medicine, Vanderbilt University Medical Center, Nashville, TN, USA; Vanderbilt Institute of Infection, Immunity, and Inflammation (VI4), Vanderbilt University Medical Center, Nashville, TN, USA

**Keywords:** microvilli, brush border, IMAC, CDHR2, CDHR5, intermicrovillar adhesion, ileum, mucosal bacteria, segmented filamentous bacteria

## Abstract

Enterocytes present a continuous array of uniform, tightly packed, apical microvilli known as the “brush border”. Adjacent microvilli tips are linked by an intermicrovillar adhesion complex (IMAC) composed of cadherins that are required for microvillar packing, length uniformity, and junctional integrity, with disruption linked to intestinal autoimmune disease and infections. We found that IMAC-deficient mice have dysbiosis with increased adherent (actin-dependent) and mucosal (actin-independent) bacteria in the terminal ileum. The bacteria are primarily commensals without differences in diversity between groups. Primary segments of Segmented filamentous bacteria (SFB) recruit the microvilli-associated proteins EPS8 and IRTKS, as well as the Arp2/3 complex, to remodel actin. SFB not only anchor adjacent to epithelial cell junctions but also incorporate ZO-1 into their attachment structures. Together, these data reveal maturation of the SFB-host interface that mirrors the complex SFB lifecycle. Our work indicates a previously unrecognized role for the brush border as a key component of host defense against luminal microbes.

## INTRODUCTION

The small intestine is essential for life, serving as the sole site of nutrient absorption. The epithelial lining is composed of a continuous layer of cells, including enterocytes, goblet cells, tuft cells, and enteroendocrine cells, with enterocytes being the most abundant. The surface of the small intestine is organized into finger-like projections called villi, while individual enterocytes display tightly packed apical membrane protrusions called microvilli, which optimize nutrient absorption by increasing surface area. The apical membrane of each enterocyte contains thousands of uniform microvilli, collectively known as the brush border. Intermicrovillar adhesion complexes (IMACs) link the tips of adjacent microvilli to maintain their dense packing and consistent length^1, 2^.

Disruption of IMACs has been implicated in intestinal inflammatory disease as a potential precursor indicative of underlying epithelial defects or as a response to chronic inflammation, damage, or dysbiosis^3–7^. Understanding how IMAC dysfunction shapes the mucosal environment at the epithelial surface will help clarify fundamental mechanisms of intestinal homeostasis and disease.

The small intestine acts as a defensive barrier against microbes and toxins in the intestinal lumen^8^. The brush border is the first point of contact between the intestinal epithelium and luminal microbes, with its dense microvilli preventing bacterial attachment and internalization^9^. Numerous pathogenic bacteria remodel the actin cytoskeleton to exploit host cells for attachment and reproduction, including *Listeria monocytogenes*^10^, enteropathogenic *Escherichia coli* (EPEC)^11^, enterohemorrhagic *Escherichia coli* (EHEC)^11^, and *Salmonella typhimurium*^12^. While pathogen-host interactions have been extensively studied, far less is known about commensal interactions with the epithelial surface in the ileum as they typically live in suspension without directly contacting the epithelium^13^.

Microvilli consist of a membrane-wrapped cores of parallel actin filaments (F-actin) which extend into the sub-apical terminal web where microvilli are anchored^14^. The distal tips of actin cores are embedded in an electron-dense tip complex containing Epidermal growth factor pathway substrate 8 (EPS8) and insulin receptor tyrosine kinase substrate (IRTKS)^15–17^. These proteins are essential for initiating microvillus growth and maintaining the stability of immature microvilli^16^. The F-actin core is organized by actin-bundling proteins villin-1 (VIL1)^18^, Espin (ESPN)^19^, Fimbrin (also known as Plastin, PLS1)^20^, and Mitotic interactor and substrate of PLK1 (MISP)^21^. IMACs bridge adjacent microvilli through extracellular linkages formed by trans-heterophilic interactions between Cadherin-Related Family Proteins CDHR2 and CDHR5^2^. The CDHR2/CDHR5 intracellular domains connect to the F-actin core via protein scaffolds and a myosin motor/light chain that transports the complex microvilli tips (Figure 1A) ^2, 22–27^. Loss of any one IMAC component disrupts adhesion and prevents tip localization.

**Figure 1.**
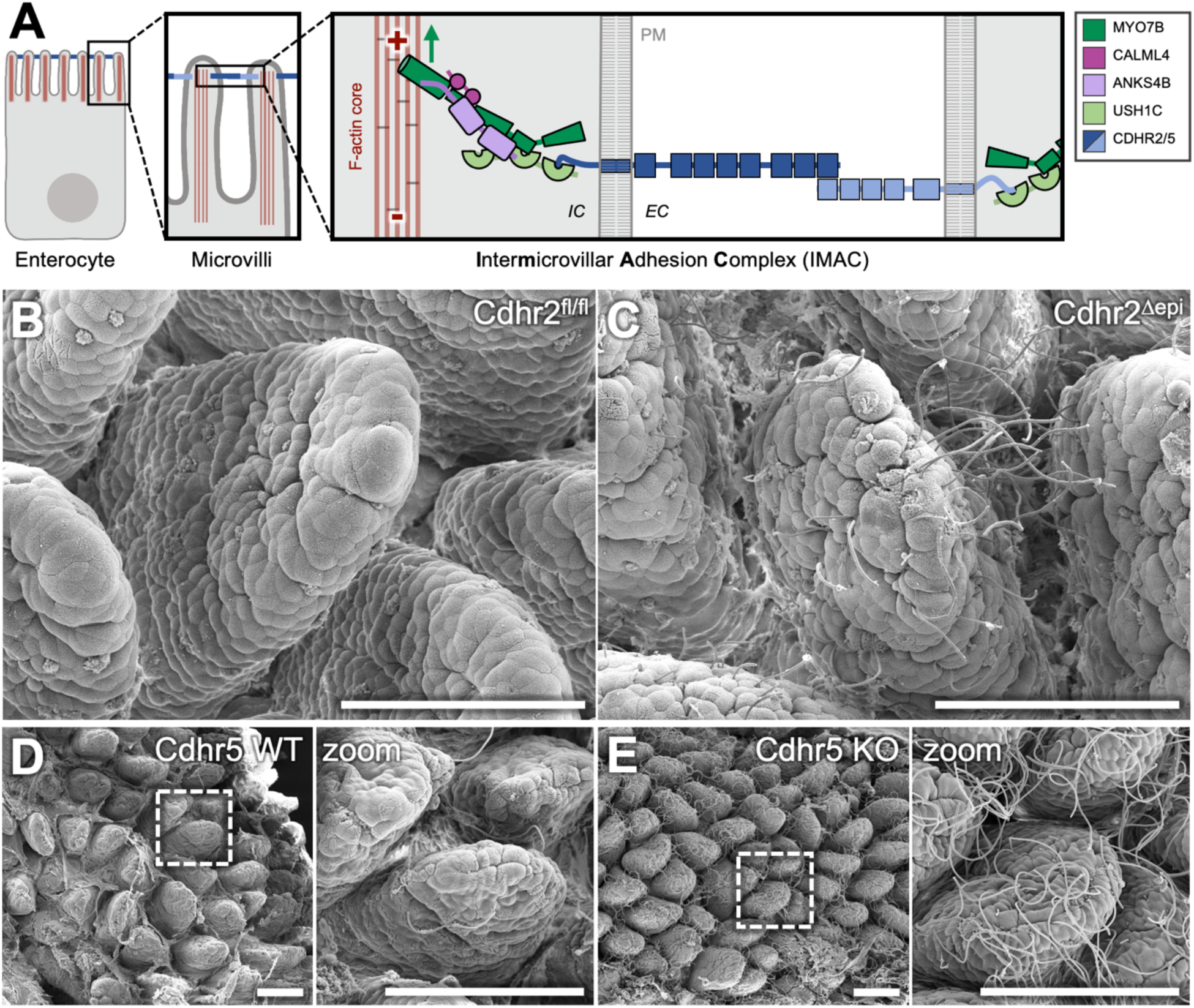
Mice with a disrupted IMAC have increased adherent bacteria in the terminal ileum. (A) Diagram highlighting components of the intermicrovillar adhesion complex (IMAC) at the tips of microvilli. PM = plasma membrane, IC = intracellular, EC = extracellular. Red “+” and “-” indicate F-actin barbed and pointed ends, respectively. Green arrow indicates F-actin barbed-end directed motion of MYO7B. (B) Representative SEM image of Cdhr2^fl/fl^ mouse terminal ileum. (C) Representative SEM image of Cdhr2^Δepi^ mouse terminal ileum. (D) Representative SEM image of Cdhr5 WT mouse terminal ileum. Boxed area enlarged in zoom. (E) Representative SEM image of Cdhr5 KO mouse terminal ileum. Boxed area enlarged in zoom. Scale bars 100 µm (B-E).

IMACs also connect microvilli across adjacent enterocytes, creating a continuous luminal brush border^28^. Disruption of IMACs results in junctional defects, including reduced localization of junctional proteins, loss of apical tension, and decreased transepithelial electrical resistance (TEER)^28–30^. Despite IMAC disruption, brush border formation remains robust with significant functional redundancy. Notably, a triple knockout of microvillus bundling proteins (VIL1, ESPN, PLS1) remains viable, retaining the ability to form microvilli^31^. Knockout mice with defects in microvilli ultrastructure and organization are viable and relatively healthy at baseline with deficits (tissue fragility^32^, increased sensitivity to DSS^32–34^) revealed only under stress. Mice lacking the IMAC component, CDHR5, exhibit increased sensitivity to DSS, including higher levels of invasive bacteria^34^.

Defects in microvilli structure have been linked to intestinal autoimmune disease including Celiac Disease^3^, Crohn’s Disease^4^ and Ulcerative Colitis^34^. Genes critical for microvilli organization, including IMAC components, are downregulated in intestinal samples from patients with Crohn’s Disease, even in the absence of active inflammation^4^. Additionally, shortened microvilli have been proposed as a biomarker predicting non-response to biologic therapies^7^. Single cell RNA sequencing from patients with Ulcerative Colitis demonstrated CDHR5 is downregulated in inflamed areas^34^. These findings implicate IMAC disruption in chronic intestinal inflammatory diseases, suggesting IMAC disruption is a marker of underlying epithelial defects or occurs in response to dysbiosis, damage or inflammation.

In this study, we report that IMAC-deficient mice develop small intestine dysbiosis. Both actin-dependent (adherent) and actin-independent (mucosal-associated) bacteria make direct contact with the epithelial surface of the terminal ileum not associated with shifts in microbiota composition. We identify the actin-dependent bacteria as Segmented filamentous bacteria (SFB). SFB colonize the terminal ileum of IMAC-deficient mice at a high levels, recruiting the microvilli-associated proteins EPS8 and IRTKS, as well as the Arp2/3 complex, to remodel the actin cytoskeleton. SFB anchor adjacent to epithelial junctions and incorporate junctional protein ZO-1 into their attachment structures. These findings reveal IMAC-mediated organization of the brush border as a key component of host defense against luminal microbes. The loss of separation between luminal bacteria and the epithelial surface provides in IMAC-deficient mice provides a model system for studying host-microbe interactions and the role of epithelial morphology in dysbiosis.

## Results

### Mice with a disrupted IMAC have increased adherent bacteria in the terminal ileum

To understand how disruptions in microvilli packing and organization alter small intestinal physiology *in vivo*, we examined the terminal ileum of mice with and without tissue-specific disruption of intermicrovillar adhesion. *Cdhr2^fl/fl^* mice express full-length Cdhr2 but harbor a conditional deletion allele that, when crossed with villin-Cre expressing mice, results in tissue-specific deletion of exons 4 and 5, leading to a deletion of Cdhr2 in intestinal epithelium referred to herein as *Cdhr2^Δepi^* (Figure S1A and previously described^30^). Whole mount scanning electron microscopy (SEM) of the terminal ileum revealed a striking population of filamentous bacteria adherent to the epithelial surface in *Cdhr2^Δepi^* mice, with bacteria concentrated distally on villi (Figure 1C). By contrast, *Cdhr2^fl/fl^* control mice had few if any visible adherent bacterial filaments (Figure 1B). To determine if the increase in adherent bacteria was specific to *Cdhr2^Δepi^* mice, we examined terminal ileum tissue from mice lacking an alternate IMAC component, Cdhr5. We again observed a dramatic increase in adherent filamentous bacteria in the ileum of Cdhr5 KO mice compared to WT control mice (Figure 1D-E). Cdhr5 WT mice had minimal adherent filamentous bacteria (Figure 1D). Based on similar results from two separate KO mice that disrupt intermicrovillar adhesion and brush border organization, we conclude that loss of IMAC function, due to loss of Cdhr2/5, facilitates bacterial adhesion to the epithelial surface and augmented colonization of the terminal ileum. Moving forward, our studies focused on the *Cdhr2^fl/fl^*and *Cdhr2^Δepi^* mice as the tissue specific deletion allows us to focus on the role of the intestinal epithelium.

### Mice with a disrupted IMAC have multiple populations of adherent bacteria enriched in the terminal ileum

The SEM whole mount studies (Figure 1B-E), while qualitatively dramatic, posed a quantification challenge due to the dense and overlapping bacteria, as well as non-uniformity in visualizing bacterial attachments in tissue. To quantify adherent bacteria, we used confocal microscopy of whole mount tissue. DAPI was used to mark epithelial cell nuclear DNA and bacterial DNA (Figure 2A), then segmented and viewed in separate channels (Figure 2B). Tissue was co-stained with phalloidin (F-actin), highlighting the sites of bacterial attachment to the brush border (Figure 2C, white arrowheads). *Cdhr2^Δepi^* mouse ileum contained significantly more bacterial attachments (Figure 2G). The F-actin attachments colocalized with chains of bacterial DNA, consistent with the filaments seen by SEM. However, confocal imaging revealed an additional population of adherent single bacteria (Figure 2D), which were also enriched in *Cdhr2^Δepi^* mouse ileum (Figure 2I). Given that adherent bacteria often target and disrupt microvilli^35–38^, we assessed the localization of brush border actin-binding proteins, to characterize the adherent bacteria. EPS8 is a microvillus tip protein that binds actin barbed ends and is essential for microvilli formation and stability^16, 17, 39, 40^. In *Cdhr2^fl/fl^* and *Cdhr2^Δepi^* mice, EPS8 localized to only a subset of adherent bacteria (Figure 2E-F). While present in both *Cdhr2^fl/fl^* and *Cdhr2^Δepi^* mice, EPS8-containing attachments were significantly increased in *Cdhr2^Δepi^*mice (Figure 2H) particularly within single bacteria (Figure 2J). However, most adherent bacteria were filamentous and lacked EPS8 staining (Figure 2G, K).

**Figure 2.**
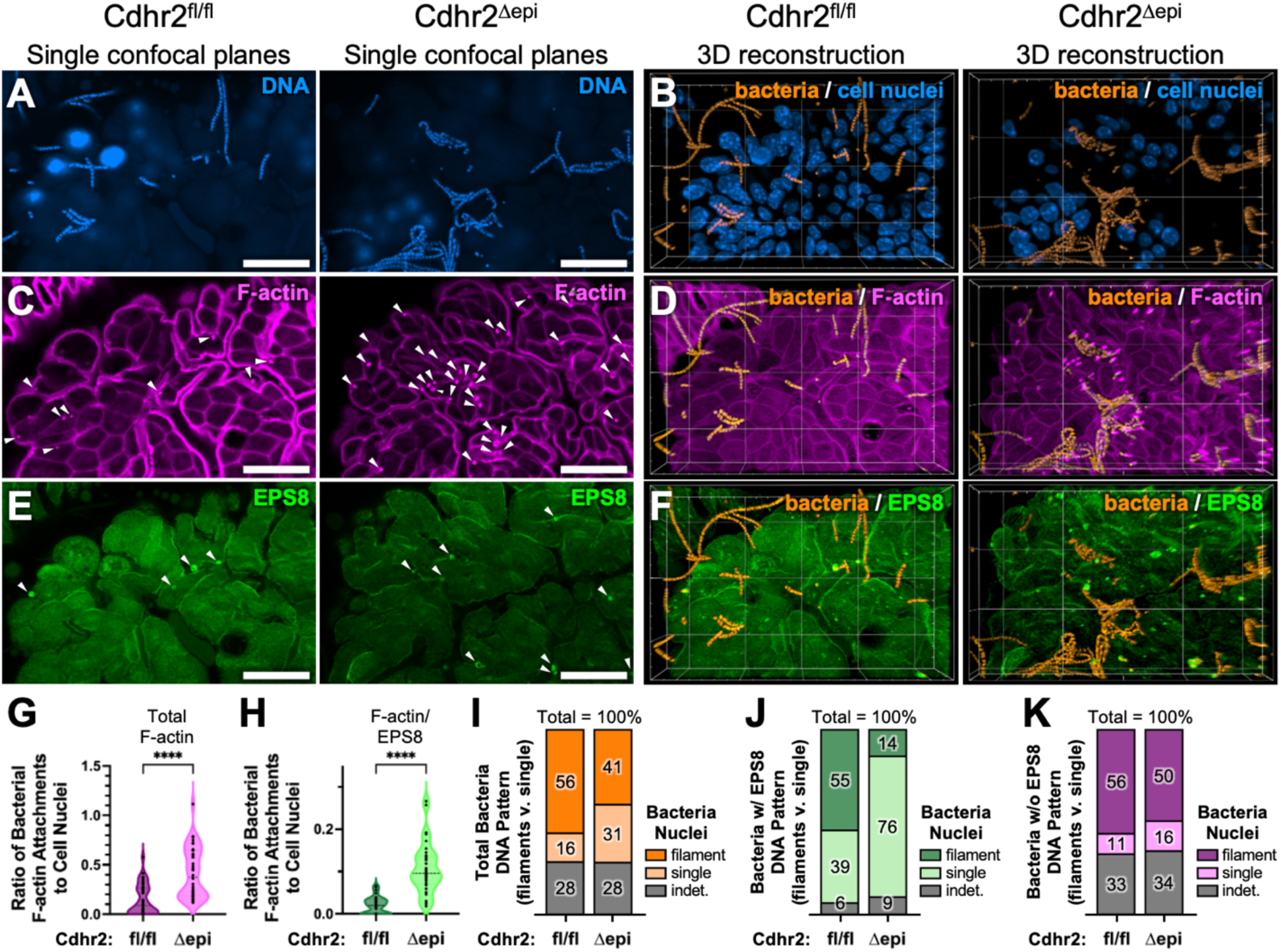
Mice with disrupted intermicrovillar adhesion have multiple populations of adherent bacteria enriched in the terminal ileum. Whole mount confocal images of ileum tissue comparing Cdhr2^fl/fl^ and Cdhr2^Δepi^ littermates. (A) Single confocal plane stained with DAPI to mark DNA. (B) 3D rendering of DAPI segmented using an artificial intelligence (AI) algorithm to isolate and differentially contrast bacterial DNA (orange) compared to cell nuclei (blue). (C) Single confocal plane stained with phalloidin to mark F-actin (actin-dependent adherent bacteria, white arrowheads). (D) 3D rendering of AI segmented bacterial DNA (bacteria) and phalloidin (F-actin). (E) Single confocal plane stained for EPS8 (actin-dependent adherent bacteria with EPS8, white arrowheads). (F) 3D rendering of AI segmented bacterial DNA (bacteria) and EPS8. (G-H) Quantification of actin-dependent adherent bacteria (Total F-actin, G) or actin-dependent adherent bacteria with EPS8 (F-actin/EPS8, H) relative to epithelial cell nuclei on distal villi. Each data point represents counts from one villus. For G, n = 39 Cdhr2^fl/fl^ villi (765 bacterial attachments / 5921 cell nuclei) and n = 40 Cdhr2^Δepi^ villi (2389 attachments / 5657 cell nuclei) taken from 5 sets of paired littermates. For H, n = 33 Cdhr2^fl/fl^ villi (95 bacterial attachments / 5134 cell nuclei) and n = 33 Cdhr2^Δepi^ villi (456 attachments / 4469 cell nuclei) taken from 4 sets of paired littermates. Violin plots with a dotted line at the median. Significance calculated using the Mann-Whitney test, **** p < 0.0001. Median 0.085 v. 0.377 actin-dependent bacterial attachments/epithelial cell in Cdhr2^fl/fl^ and Cdhr2^Δepi^ villi respectively. Median 0.022 v. 0.096 actin-dependent bacterial attachments with EPS8/epithelial cell in Cdhr2^fl/fl^ and Cdhr2^Δepi^ villi respectively. (I-K) Stacked bar charts of adherent bacteria utilizing F-actin attachments by bacterial DNA morphology (filament, single, or indeterminate) total F-actin adherent bacteria (I) total F-actin adherent bacteria with EPS8 (J) and F-actin adherent bacteria without EPS8 (K). Scale bars 20 µm (A,C,E), and background grid 20 µm (B,D,F).

Given the increased level of adherent bacteria in *Cdhr2^Δepi^*mice, we evaluated the ileum for signs of inflammation. Hematoxylin and eosin (H&E) staining of the small intestine did not reveal evidence of tissue damage or increased immune infiltrates (data not shown). We used qRT-PCR to quantify the expression of inflammatory genes commonly elevated in bacterial infections including the chemokine *Cxcl1*, cytokines (*Il22, Il17a,* and *Il6)*, and the bactericidal antimicrobial peptide *Reg3g*. There were no differences between *Cdhr2^fl/fl^* and *Cdhr2^Δepi^*mice (Figure S1C). We also performed flow cytometry to phenotype the lymphocytes in the lamina propria of the small intestine (Figure S1F), mesenteric lymph nodes (data not shown), and inguinal lymph nodes (data not shown) from age- and sex-matched *Cdhr2^Δepi^* and *Cdhr2^fl/fl^*littermates. Representative plots of the gating scheme used to identify T cell subsets and IL-17A and IFN-γ expressing cells are shown (Figure S1D and E, respectively). T cell subsets (CD4, CD8, and gamma-delta (ψο)) were not significantly different in the lamina propria (Figure S1F). Further, the percentage of interferon-gamma (IFN-γ) and IL-17A-producing cells in these T cell subsets did not differ (Figure S3G). There was a trend towards higher IL-17A expression in ψο T cells in the mesenteric lymph nodes of *Cdhr2^Δepi^*mice but this was not significant (data not shown). Overall, this indicates that while *Cdhr2^Δepi^* mice have higher numbers of bacteria in contact with the intestinal epithelium including formation of F-actin based attachments, the mice are not mounting an enhanced systemic or tissue-resident inflammatory response.

### Loss of intermicrovillar adhesion facilitates polymicrobial tissue colonization with bacteria utilizing actin-dependent and actin-independent surface attachments

To understand if loss of intermicrovillar adhesion results in colonization with specific bacteria, we sought to identify the ileum-associated bacteria in *Cdhr2^fl/fl^*and *Cdhr2^Δepi^* mice. The filamentous bacteria morphology observed by SEM and whole mount imaging are suggestive of *Candidatus Arthromitis*, also known as segmented filamentous bacteria (SFB) which preferentially colonize the mouse ileum shortly after weaning then decline to very low levels over several weeks^41, 42^. We performed fluorescence *in situ* hybridization (FISH) directed against SFB 16s rRNA (SFB FISH)^43^. Filamentous bacteria on sectioned ileum from *Cdhr2^Δepi^* mice stained with the SFB FISH probe (Figure 3A). While whole mount *Cdhr2^fl/fl^* mouse ileum demonstrated low levels of adherent bacteria, the low density of adherent bacteria consistently resulted in sectioned tissue with no tissue-associated bacteria limiting analysis (data not shown). Staining *Cdhr2^Δepi^*ileum for actin, DNA, and SFB FISH confirmed SFB directly contacts the F-actin rich brush border (Figure 3B, zoom 1). Together these data establish that the adherent filaments in *Cdhr2^Δepi^* mice are SFB.

**Figure 3.**
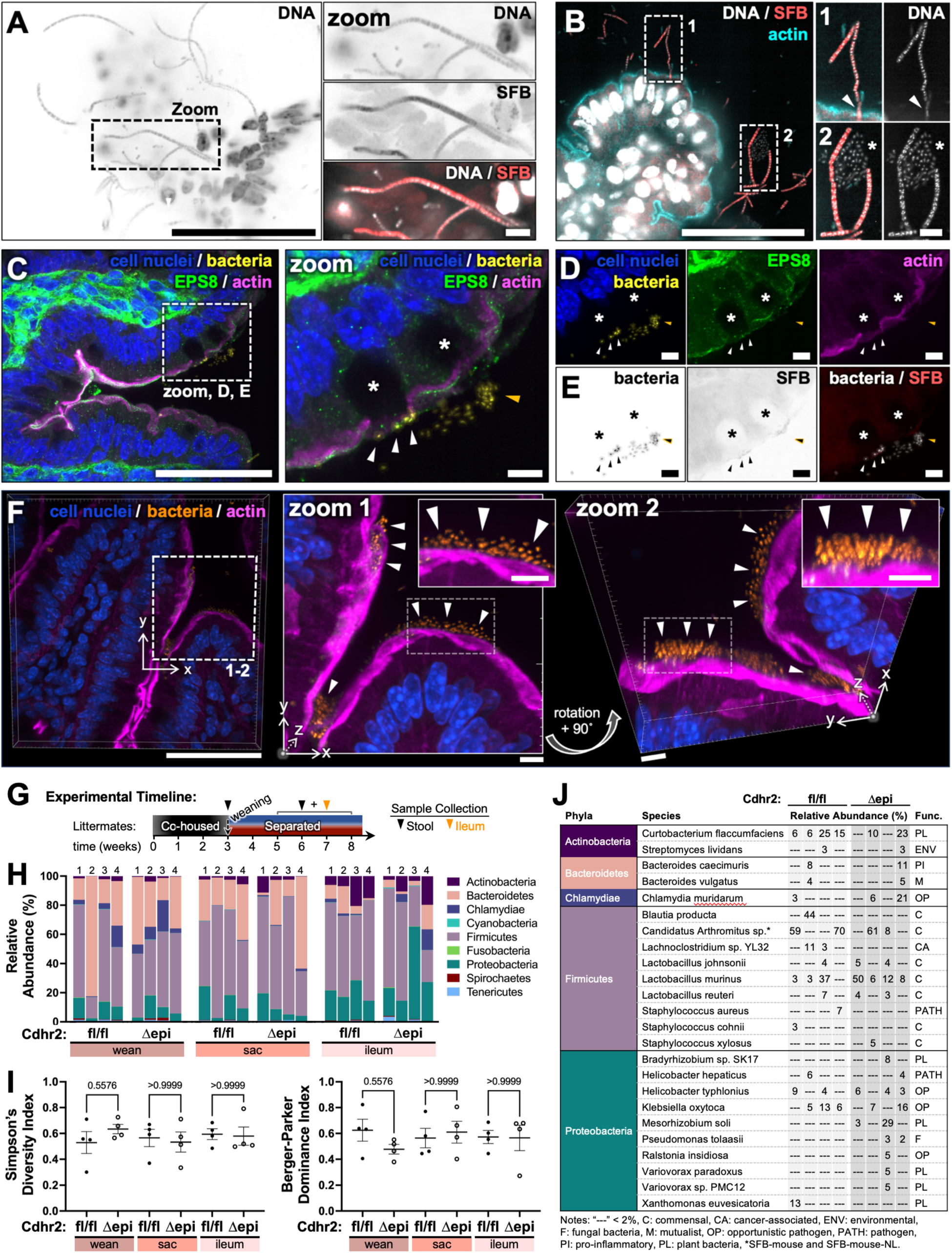
Loss of intermicrovillar adhesion facilitates polymicrobial tissue colonization with bacteria utilizing actin-dependent and actin-independent surface attachments. (A) Single confocal slice of Cdhr2^Δepi^ ileum frozen section stained for DNA (Hoechst) and SFB 16S-rRNA fluorescence *in situ* hybridization (SFB FISH). Boxed area enlarged in zoom. (B) Deconvolved single confocal slice of paraffin sectioned Cdhr2^Δepi^ mouse ileum stained for DNA (DAPI), SFB 16S-rRNA FISH (SFB), and actin. Boxed areas enlarged in Zooms 1 and 2 contrast adjusted to visualize bacteria. (Zoom 1-2) left images show a 3-color overlay DNA (white), SFB (red), actin (cyan), and right images with DNA only. White arrowhead highlights bacteria interaction with the surface actin/brush border, and the asterisk marks a cluster of bacteria negative for SFB 16S-rRNA FISH. (C) Maximum intensity projection (maxIP, depth 11.25 µm) of paraffin sectioned Cdhr2^Δepi^ mouse ileum stained for DNA (DAPI), EPS8, and actin. DAPI segmented using an AI algorithm to isolate and differentially contrast bacteria (yellow) relative to cell nuclei (blue). Boxed area enlarged in Zoom, D, and E. Asterisks mark goblet cells, arrowheads highlight bacteria associated with goblet cells/mucus. (D) Individual channels from C, segmented DAPI (cell nuclei/bacteria), EPS8, and actin. (E) Individual channels from C inverted and shown in grayscale, segmented DAPI (bacteria), SFB 16S-rRNA FISH (SFB), and overlay. (F) 3D rendered maxIP (depth 8.40 µm) of paraffin sectioned Cdhr2^Δepi^ ileum stained for DNA (DAPI) and actin. DAPI is AI segmented to isolate and differentially contrast bacteria (orange) relative to cell nuclei (blue). Boxed areas—enlarged and contrast enhanced (zoom 1); enlarged, contrast enhanced and rotated (zoom 2)—to visualize tissue-associated bacteria. Boxed areas in zoom images further enlarged and contrasted in insets, upper right. White arrowheads indicate bacteria. (G) Diagram of the experimental timeline. Paired Cdhr2^fl/fl^ and Cdhr2^Δepi^ littermates are co-housed until weaning. Stool collected at weaning (∼3 weeks of age), then mice are housed separately by genotype. Stool and ileum tissue are collected between 5-8 weeks of age. (H) Relative abundance of bacterial phyla shown as stacked bar graphs. Samples collected from Cdhr2^fl/fl^ and Cdhr2^Δepi^ mice (sets 1-4); Stool at Wean (Wean), Stool at sacrifice (Sac), ileum tissue at sacrifice (Ileum). (I) Bacteria phylum alpha diversity. Simpson’s Diversity Index (left) and Berger-Parker Dominance Index (right). Error bars represent mean ± SEM. Significance calculated using the Friedman test. (J) Table of bacteria species relative abundance detected > 2%. Scale bars: 50 µm (A), 5 µm (A, zoom), 50 µm (B), 5 µm (B, zoom 1-2), 50 µm (C), 5 µm (C, zoom), 5 µm (D), 5 µm (E), 50 µm (F), 5 µm (F, zooms 1-2 and insets).

Non-SFB bacteria were also identified in contact with the epithelial surface, but they lacked F-actin attachments. We identified clusters of bacteria and scattered individual bacteria consistently associated with SFB (Figure 3B, zoom 2; Figure 4F, zoom 3). Volume reconstruction demonstrated bacteria associated with villus goblet cells (Figure 3C). Actin staining delineates the brush border, and EPS8 marks the distal tips of microvilli. Goblet cells are visible as gaps in the apical brush border associated with a cytoplasmic void where mucus is stored or was recently secreted (Figure 3C, Figure S2A). Immunofluorescence (IF) and EM tissue processing preserves apical mucus associated with goblet cells as seen on SEM (Figure S2B). Bacteria are observed in direct contact with goblet cells—typically at the cell periphery (Figure 3C zoom, white arrowheads) or in the adjacent lumen—suggestive of mucus binding (Figure 3C zoom, orange arrowheads). An oblique view more clearly highlights three goblet cells with associated bacteria (Figure S2C). Viewed as separate channels, bacteria are again seen at the goblet cell surface (Figure 3D, Figure S2D). These bacteria do not stain with the SFB FISH probe (Figure 3E). Additionally, tissue-associated bacteria were seen in large groups overlying the epithelial surface (Figure 3F).

**Figure 4.**
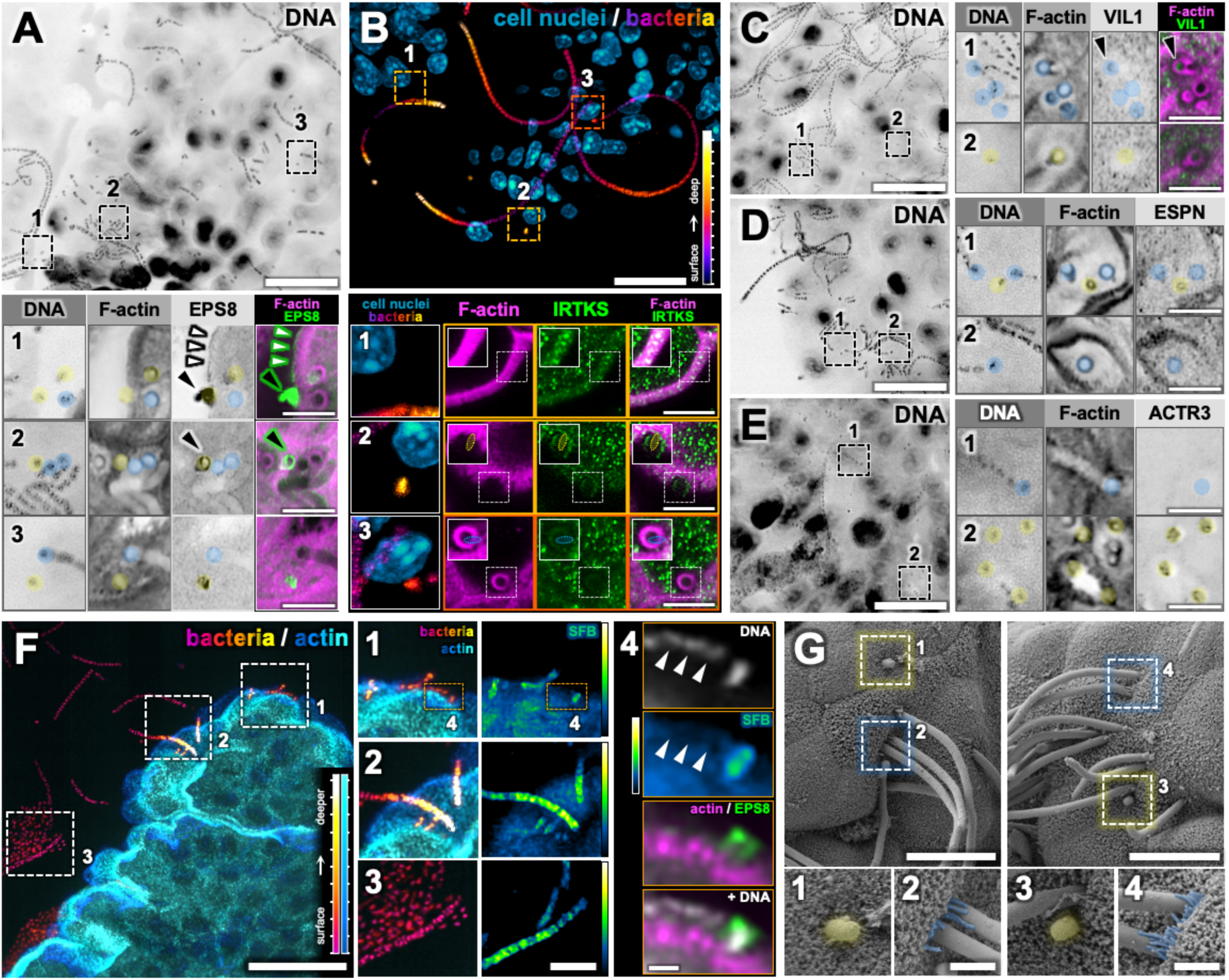
Adherent primary SFB segments recruit microvilli proteins (EPS8 and IRTKS) and remodel the actin cytoskeleton via Arp2/3. (A) Single confocal plane at the villus surface of Cdhr2^Δepi^ mouse ileum whole mount tissue stained with DAPI (DNA). Boxed areas enlarged (zooms 1-3) with additional stains DAPI (DNA), phalloidin (F-actin), EPS8, and F-actin/EPS8 overlay. Adherent bacteria marked with a transparent overlay (yellow, single bacteria; blue, filaments). White arrowheads highlight EPS8 localization in enterocytes (zoom 1). Teardrop-shaped bacteria indicated by black arrowheads (zooms 1-2). (B) Cdhr2^Δepi^ mouse ileum frozen tissue section stained with DAPI (DNA), deconvolved, AI segmented to isolate and differentially contrast bacterial DNA and viewed as a maxIP (depth 12.6 µm). The bacteria channel is z-depth color-coded; closer bacteria purple/magenta, deeper bacteria yellow/white, z-depth color-code at right. DNA from cell nuclei (cyan/blue). Boxed areas enlarged below (zooms 1-3) with the box color indicating the z-depth of the zoom images. (B, zoom 1) IRTKS localization in the enterocyte brush border. (B, zooms 2-3) highlights adherent bacteria. Column 1 is the main image panel enlarged (maxIP, depth 12.6 µm). Subsequent columns 2-4 show additional stains phalloidin (F-actin), IRTKS, and F-actin/IRTKS overlay as a maxIP (depth 0.3 µm). White boxed areas within zoom images (columns 2-4) are contrast enhanced and shown as insets (upper left) with bacteria localization indicated by ovals (yellow, single bacteria; blue, filaments). (C-E) Single confocal planes at the villus surface of Cdhr2^Δepi^ mouse ileum whole mount tissue stained with DAPI (DNA). Boxed areas (zooms 1-2) enlarged at right. DAPI (DNA) and phalloidin (F-actin) are shown in all sets plus Villin (VIL1, C), F-actin/VIL1 (C), Espin (ESPN, D), and Arp3 (ACRT3, E). (C, zoom 1), black arrowheads indicate a filamentous attachment with eccentric VIL1 localization. (F) Cdhr2^Δepi^ mouse ileum paraffin tissue section stained for DAPI (DNA) and actin, deconvolved, and viewed as a maxIP (depth 10.4 µm). A z-depth color-code is applied to each channel, scales lower right. DNA AI segmented to isolate bacteria, cell nuclei are not displayed. Boxed areas enlarged at right (zooms 1-3). Column 1, bacteria/actin maxIP (depth 10.4 µm) enlarged from the main panel. Column 2, intensity color-coded SFB FISH (SFB) maxIP (depth 10.4 µm), intensity scale at right). Boxed areas in zoom 1 further enlarged in zoom 4. Z-depth indicated by box color, based on the main panel bacteria z-depth color-code. (F, zoom 4) Single confocal plane of surface bacteria with additional stains; DAPI (DNA) without AI segmentation, SFB FISH (SFB) intensity color-coded, actin/EPS8 overlay, and actin/EPS8 plus DNA overlay. White arrowheads indicate SFB FISH negative bacteria. (G) SEM images of Cdhr2^Δepi^ mouse terminal ileum. Boxed areas enlarged in zooms 1-4 to highlight adherent bacteria (yellow, single bacteria; blue, filaments). (G, zooms 1, 3) single bacteria. (G, zooms 2, 4), filamentous bacteria with elongated microvilli at the base. Scale bars: 20 µm (A-F, main panels), (B, F) z-depth color-code tick marks at 1 µm intervals, 5 µm (A-F zooms 1-3), 2.9 µm (B, zoom 1-3, insets), 1 µm (F, zoom 4), 10 µm (G, main panel), 2 µm (G, zooms 1-4).

Zoom images reveal layered colonies of bacteria (white arrowheads). The bacteria do not utilize F-actin attachments; but rather are closely apposed to the brush border, mirroring but not disrupting the surface F-actin (Figure S3A). 3D surfaces generated from the confocal data demonstrate the relationship between bacteria and the brush border in Cdhr2^Δepi^ mice. (Figure S3B-C). The bacteria establish a complex, multi-layered structure (at least 25 x 4 µm in xy and 4 µm in z, Figure S3B), consistent with biofilm formation (Figure 3F, zoom 1-2). Rotational views highlight the 3D structure of the bacterial colony (Figure S3B). Moreover, the bacteria consistently lie above the actin which maintains a smooth, continuous surface (Figure S3C). Like goblet cell-associated bacteria, these bacteria do not stain with the SFB FISH probe (Figure S3D). Together, these data show IMAC disruption allows multiple populations of bacteria to colonize the epithelial surface. Moreover, bacteria employ multiple attachment methods including anchoring to epithelial cells utilizing F-actin, colonization of adherent bacteria, mucus-binding, and surface colony/biofilm formation.

While the adherent filamentous bacteria colonizing the terminal ileum are identified as SFB, the other mucosal-associated bacteria remain unknown. We used shotgun metagenomic sequencing to identify mucosal bacteria and to assess for microbial shifts in *Cdhr2^Δepi^* mice compared to *Cdhr2^fl/fl^* mice. Paired sets of littermates were initially co-housed to establish consistent maternal microbial exposure, then separated by genotype at weaning (Figure 3G). Stool was collected immediately prior to separation, and stool and ileum tissue were collected between 5 and 8 weeks of age. Relative abundance of bacterial phyla shown as stacked bar graphs (Figure 3H). There were no significant differences in alpha diversity using either the Simpson’s Diversity Index or the Berger-Parker Dominance Index (Figure 3I). Beta diversity calculations were not completed due to the high level of variation in read count. These data confirm polymicrobial bacteria associate with ileum tissue, dominated by commensals, but opportunistic pathogens, pro-inflammatory bacteria, and potential pathogens were also detected (Figure 3J). Plant bacteria were thought to be due to residual food/luminal contents. Environmental bacteria were detected in only a few samples and considered insignificant. The most abundant bacteria identified in both groups of mice were SFB and *Lactobacillus murinus*; both known to colonize the small intestine. Many commensals including *Lactobacillus*, do not adhere directly to the epithelial surface, and are thought to attach to the overlying mucus or extracellular matrix^44, 45^ consistent with the patterns seen in non-SFB bacteria at the epithelial surface (Figure 3C-F). Overall, these data demonstrate that disruption of IMAC function allows bacteria to encroach on the epithelial surface without markedly changing the microbiome composition.

### Adherent primary SFB segments recruit microvilli proteins (EPS8 and IRTKS) and remodel the actin cytoskeleton via Arp2/3

Given that brush border microvilli are the first point of epithelial cell contact for luminal bacteria, we hypothesized that the bacteria colocalized with EPS8 targeted microvillar proteins to remodel brush border actin. Focusing on *Cdhr2^Δepi^*mice which have significantly more adherent bacteria compared to *Cdhr2^fl/fl^*mice, we used an immunostaining approach to define the molecular composition of the bacterial attachments. Mucosal bacteria were evident in low magnification images of DNA at the villus surface (Figure 4A-E). Z-depth color-coding highlights the 3D nature of the adherent bacteria (Figure 4B). In enterocytes, EPS8 is normally localized to the tips of microvilli (Figure 4A, zoom 1, white arrowheads). However, EPS8 also localizes to a subset of adherent bacteria (Figure 2E-F). Areas of moderate bacteria density were assessed at high magnification to allow clear identification of single versus filamentous bacteria associated with attachment sites. EPS8 colocalized only with single bacteria (Figure 4A, zooms 1-3, yellow overlay). Adjacent filaments lacked EPS8 signal (Figure 4A, zooms 1-3, blue overlay). Strikingly, the EPS8 staining at times adopts a teardrop shape (Figure 4A, zooms 1-2, black arrowheads), reminiscent of SFB holdfast segments (the first segment of SFB attached to the epithelium)^46^.

Next, we assessed the localization of IRTKS, a member of the microvillus tip complex and an EPS8 binding partner^17, 47^. IRTKS is an I-BAR (inverse-bin-amphiphysin-Rvs) protein which link changes in membrane structure with actin remodeling^48, 49^. I-BAR domains sense and induce membrane curvature^47^. IRTKS also contains an SH3 domain to recruit signaling partners and a WH2 domain that binds actin monomers. Within enterocytes, IRTKS localizes to the distal tips of actively growing microvilli, recruiting EPS8, and promoting microvilli growth^16, 17, 50^. In the mature brush border IRTKS localizes along the length of microvilli (Figure 4B, zoom 1)^16, 17, 50^. We identified IRTKS enriched at the surface of single adherent bacteria, best visualized when captured in cross section in the xy-imaging plane (Figure 4B, zoom 2, inset). Adherent SFB filaments contained faint IRTKS colocalized with the F-actin attachment site, (Figure 4B, zoom 3, inset) and more prominently in a ring where the embedded holdfast segment touches the cell surface, but lacked the surface localization seen in single bacteria. These data demonstrate the EPS8-positive adherent bacteria recruit microvilli tip proteins associated with actin remodeling to their sites of attachment.

Given adherent single bacteria recruit microvillar tip proteins (EPS8/IRTKS), we sought to determine whether the bacteria directly incorporate F-actin from microvilli into their attachment sites. Cellular arrays of F-actin are made parallel bundles or branched filaments and can be rapidly remodeled. The actin structure is defined by actin-binding proteins and the actin nucleator used to create new filaments. We used an immunostaining approach to assess the actin structure within bacterial attachments. VIL1 and ESPN are bundling proteins that maintain the parallel F-actin bundles in microvilli^18, 19^. VIL1 was not enriched at single bacteria attachments (Figure 4C, zoom 2). VIL1 was occasionally seen at filamentous SFB attachment sites (Figure 4C, zoom 1, black arrowhead) but more consistently was not present at filamentous SFB attachments (Figure 4C, zoom 1). In contrast, ESPN consistently colocalized with F-actin attachments of both single bacteria and SFB filaments (Figure 4D, zooms 1-2). However, ESPN intensity was markedly lower than within the brush border. These results do not support bundled, parallel F-actin as the primary structure underlying bacterial attachments. Given the paucity of bundling proteins within bacterial attachment sites, we assessed markers of branched actin. The Arp2/3 complex, comprised of seven protein subunits, is the primary nucleator of branched actin^51^. We stained for ACTR3 (actin related protein 3, Arp3), a subunit of the Arp2/3 complex.

ACTR3 localized to adherent single bacteria but not to SFB filaments (Figure 4E, zooms 1-2), mirroring the pattern seen with EPS8. Based on these data, we conclude that adherent single bacteria recruit EPS8/IRTKS to facilitate actin remodeling as the presence of Arp2/3 at single adherent bacteria indicates that the bacteria are actively remodeling the actin cytoskeleton. Higher magnification SEM images of *Cdhr2^Δepi^* ileum show single adherent bacteria mixed with filamentous bacteria (Figure 4G). Of note, elongated microvilli were seen at the base of some SFB filaments which could account for the inconsistent detection of VIL1 at the edge of SFB filament attachments (Figure 4G, zooms 1-4).

EPS8 and Arp2/3 consistently localized to single bacteria with actin-dependent attachment sites while SFB filaments lacked both markers. Additionally, EPS8 positive bacteria bore striking morphological similarity to primary SFB segments, e.g. teardrop shape that tapers to a point in the attachment. Given this, we hypothesized adherent EPS8 positive bacteria represented an early step in SFB attachment (e.g. attachment of the holdfast segment). Paraffin-embedded tissue sections were co-stained for DAPI, SFB FISH, actin, and EPS8. MaxIP with z-depth color-coding affirmed the presence of multiple populations of ileum-associated bacteria (Figure 4F). Single bacteria and filamentous bacteria contact the epithelial surface (Figure 4F, zoom 1).

Filamentous bacteria are visualized nearing the brush border but are transected by the sectioning plane before contacting the cell surface (Figure 4F, zoom 2). Finally, groups of bacteria are clustered around filamentous bacteria (Figure 4F, zoom 3). SFB FISH colocalized with filamentous bacteria including short chains (Figure 4F, zooms 1-3), but also with a single bacterium at the cell surface (Figure 4F, zooms 1 and 4). Adjacent bacteria at the cell surface do not colocalize with SFB FISH (Figure 4F, zoom 4, white arrowheads), nor do the bacteria clustered around the SFB filament (Figure 4F, zoom 3) which serve as internal controls. The single SFB is in contact with the F-actin surface and colocalizes with EPS8 (Figure 4F, zoom 4), consistent with the previously identified population of EPS8 positive adherent bacteria. These data indicate SFB primary segments recruit EPS8/IRTKS and Arp2/3 to remodel surface actin generating their distinctive actin-dependent “holdfast” points. The mature SFB filaments lack these molecular markers of actin remodeling.

### SFB actin-dependent attachments incorporate ZO-1 in continuity with junctional complexes

Previous studies established CDHR2 knockout results in cell-cell junction defects including reduced localization of junctional proteins (including ZO-1) and decreased MYH4, suggestive of lower junctional tension^29, 30^. *Cdhr2^Δepi^*mice have abnormal tissue morphology with uneven epithelial surfaces and apically “domed” enterocytes^30^. Minimal differences are apparent at low magnification (Figure 5A), but higher magnification reveals defects. Structural defects are visible throughout the villus but are most pronounced at villus tips (Figure 5A, zooms 1-4), including brush border discontinuity at cell-cell junctions (Figure 5B).

**Figure 5.**
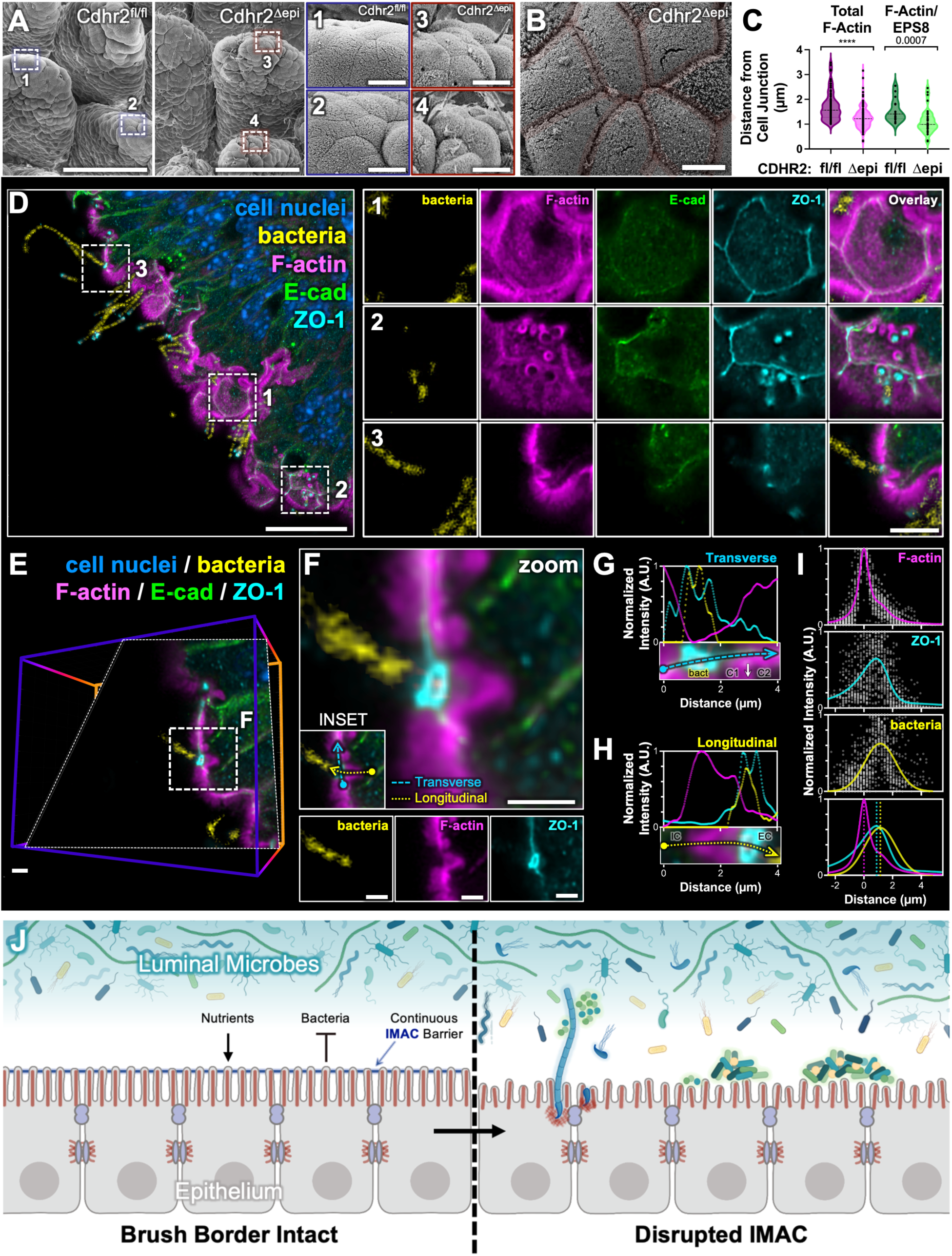
SFB actin-dependent attachments incorporate ZO-1 and are continuous with cell-cell junctions. (A) Whole mount SEM of Cdhr2^fl/fl^ and Cdhr2^Δepi^ ileum villi, boxed areas enlarged in zooms 1-4. Zooms highlight tissue morphology at the villus tips. (B) SEM of Cdhr2^Δepi^ mouse terminal ileum highlighting discontinuity of microvilli packing at cell-cell junctions (transparent red overlay). (C) Quantification of the distance from the center of actin-dependent adherent bacteria to cell-cell junctions (Total F-Actin) and the subset of actin-dependent bacteria colocalized with EPS8 (F-Actin/EPS8). Each point on the graph represents one adherent bacterium; Total F-actin Cdhr2^fl/fl^ n = 87, Cdhr2^Δepi^ n = 148; F-Actin/EPS8 Cdhr2^fl/fl^ n = 15, Cdhr2^Δepi^ n = 39; representing bacteria from 4-6 mice for each genotype. Violin plots with a dotted line at the median. Significance calculated using the Mann-Whitney test, **** p < 0.0001. Median distance from the center of adherent bacteria to the nearest cell-cell junction of 1.56 µm v. 1.22 µm (Total F-actin) in Cdhr2^fl/fl^ and Cdhr2^Δepi^ villi respectively; median distance from the center of adherent bacteria to the nearest cell-cell junction 1.41 µm v. 0.99 µm (F-actin/EPS8) in Cdhr2^fl/fl^ and Cdhr2^Δepi^ villi respectively. (D) MaxIP (depth 1.6 µm) of Cdhr2^Δepi^ ileum stained with DAPI (DNA), phalloidin (F-actin), E-cadherin, and ZO-1. DNA segmented using an AI algorithm to isolate bacterial DNA (yellow) from cell nuclei (blue). Boxed areas enlarged in zooms 1-3 show individual channels (bacteria, F-actin E-cadherin, ZO-1) and the overlay. (E) 3D volume from the upper left of (D) optically sectioned through a single adherent bacterial filament (D, zoom 3). The oblique section is parallel to the bacterial filament and perpendicular to the cell surface. Boxed area enlarged in F. (F) Oblique section through a single SFB attachment. Individual channels (bacteria, F-actin, and ZO-1) shown below. INSET indicates the Transverse and Longitudinal regions of interest analyzed in G and H. (G) Transverse fluorescence intensity distributions of F-actin, ZO-1, and bacteria of a representative adherent bacterial filament. Intensities measured at the surface extending across multiple cells (C1 = cell 1, C2 = cell 2, bact = adherent bacteria). (H) Longitudinal fluorescence intensity distributions of F-actin, ZO-1, and bacteria of a representative adherent bacterial filament. Intensities measured parallel to the bacterial filament/F-actin attachment extending from intracellular (IC) to extracellular (EC). (I) Longitudinal fluorescence intensity distributions of F-actin, ZO-1, and bacteria extending from intracellular to extracellular. Peak F-actin intensity set at 0. (J) Model of the impact of IMAC disruption on epithelial surface defense against luminal bacteria. Scale bars: 100 µm (A main panel), 20 µm (D), 10 µm (A, zooms 1-4), 5 µm (B; D, zooms 1-3), 2 µm (E-F).

Given high levels of adherent SFB in Cdhr2^Δepi^ ileum and that SFB anchor preferentially near cell-cell junctions^42^, we hypothesized SFB target junctions for attachment. Using whole mount tissue, we quantified the distance from the center of adherent bacteria to the closest cell-cell junction. SFB anchored closer to cell-cell junctions on *Cdhr2^Δepi^*ileum compared to *Cdhr2^fl/fl^* ileum (Figure 5C). The subset of SFB colocalized with EPS8 (representing initial SFB segments) also anchored closer to cell-cell junctions on *Cdhr2^Δepi^* ileum compared to *Cdhr2^fl/fl^* ileum (Figure 5C).

Little is known about the molecular composition of SFB attachments beyond the utilization of F-actin^52^. Given the proximity of adherent SFB to cell-cell junctions is enhanced when intermicrovillar adhesions are disrupted via CDHR2 deletion, we sought to determine whether SFB attachments target junctional proteins. To test this idea, we used an immunostaining approach to assess the localization of junctional proteins. Frozen tissue sections were stained for DNA (DAPI), F-actin (phalloidin), ZO-1 (tight junctions), and E-cadherin (adherens junctions). Confocal imaging of *Cdhr2^Δepi^* ileum revealed ZO-1 localized to bacterial attachment sites. In enterocytes, ZO-1 encircles the lateral edge of the cell just below the brush border forming an adhesive belt between adjacent cells (Figure 5D, zoom 1). When bacterial attachments are sectioned parallel to the cell surface, the actin-rich attachment is seen as an open ring of F-actin that tapers deeper into the cell (Figure 5D, zoom 2). ZO-1 localizes inside the F-actin ring (between the bacterial DNA and F-actin) while E-cadherin is not enriched at attachment sites. When sectioned longitudinally, the bacterial attachment is seen as a defect in the brush border with a teardrop shaped F-actin attachment extending into the cell (Figure 5D, zoom 3). ZO-1 is concentrated where the bacterium meets the cell surface. An oblique section parallel to a representative bacterial filament as it meets the cell surface more clearly shows ZO-1 encircling the primary SFB segment (Figure 5E-F) with the ZO-1 surrounding the bacterium in continuity with ZO-1 in adjacent cell-cell junctions (Figure 5F-G). ZO-1 forms a layer between the surface bacterium which remains extracellular and the deeper F-actin attachment (Figure 5H).

## DISCUSSION

CDHR2 is a core component of the IMAC. The original studies characterizing *Cdhr2^Δepi^* mice revealed profound ultrastructural brush border defects including shortened, disorganized microvilli with decreased packing density and reduced localization of multiple apical markers^30^. Despite these defects, the mice appeared healthy providing an opportunity to study how brush border disruption influences host defense *in vivo*. Our investigation focused on the terminal ileum which houses the core of the intestinal immune system and contains the highest bacterial concentration in the small intestine.

We observed small intestine dysbiosis with increased adherent bacteria in the absence of stressors in both Cdhr2^Δepi^ and *Cdhr5* KO mice. Previous studies reported increased invasive bacteria in the colon of *Cdhr5* KO only after exposure to Dextran Sodium Sulfate (DSS)^34^. In DSS-colitis, increased permeability of the inner mucus layer precedes epithelial damage and inflammation, allowing luminal bacteria to access to the epithelial surface^53, 54^. In the absence of stressors, the dense colonic mucus layer appears sufficient to compensate for epithelial defects in the colon of *Cdhr5* KO mice^34^. In contrast, the small intestine mucus is less dense and loosely associated with the epithelial surface^55^, providing less robust protection. Thus, loss of IMAC tip links in the small intestine allows luminal bacteria to attach directly to the epithelial surface, resulting in more adherent bacteria in the terminal ileum of *Cdhr2^Δepi^* and *Cdhr5* KO mice even in the absence of external stressors.

Most bacteria utilizing actin-based attachments are pathogens^11, 56, 57^; however, given the lack of infectious symptoms (e.g. weight loss or diarrhea), localization in the terminal ileum, and the characteristic tear-drop shape of the actin attachments, SFB were a prime candidate for the adherent bacteria. Non-SFB commensals identified via sequencing were initially thought to originate from residual luminal bacteria due to incomplete tissue washing. However, paraffin-embedded tissue sections (undergoing less harsh washing compared to SEM or whole mount tissue preparation) revealed both SFB and non-SFB bacteria directly interacting with the epithelial surface of *Cdhr2^Δepi^* mice. Non-SFB bacteria employ diverse actin-independent mechanisms for attachment, including sub-colonization of SFB^42^, mucus binding^58–60^, and biofilm formation^61, 62^. While these bacteria are most likely commensals, validated FISH probes for most commensal species are unavailable, limiting direct visualization. The colonization of the epithelium by bacteria using diverse attachment mechanisms indicates that intermicrovillar adhesion globally contributes to maintaining the basal separation between luminal bacteria and the epithelial surface, not solely to defense against pathogens.

Under normal conditions, commensal bacteria in the ileum do not directly contact the epithelium but remain in suspension^13^. In our study, bacteria directly contacted the epithelial surface without marked changes in microbiota composition in *Cdhr2^Δepi^* mice. Similarly, prior studies of *Cdhr5* KO mice did not show microbiota composition changes compared to *Cdhr5* WT mice^34^. Bacterial access to the epithelial surface is likely due to the disruption of the IMAC, which reduces microvilli packing density by ∼30% resulting in disordered microvilli. The resulting “liquid” packing, rather than the typical tight hexagonal arrangement of brush border microvilli^30^ combined with apical doming, increases access to the lateral microvillar membrane surfaces and intermicrovillar spaces, facilitating bacterial adhesion. Unlike host defense components such as antimicrobial peptides, which influence microbial composition^43^, IMAC disruption results in dysbiosis via altered bacterial localization^63^.

Dysbiosis of the small intestine is implicated in human diseases ranging from metabolic disorders (e.g., diabetes, obesity, malnutrition)^64, 65^ to autoimmune conditions (e.g. Crohn’s disease^66–68^, celiac disase^69^). Small intestine biofilm formation has been reported predominantly in disease states^62, 70–73^. For example, in Crohn’s disease, shifts in ileal mucosal-associated bacteria have been linked to disease onset^66^, active disease versus remission^67^, and inflamed versus uninflamed tissue within the same patient^68^. Defects in microvilli organization have been implicated in Crohn’s disease^4^ and ulcerative colitis^34^, but whether these defects are causal or secondary to prior damage, inflammation, or infection remains unclear. Our data demonstrate that microvilli defects alone are sufficient to cause dysbiosis by facilitating direct bacterial access to the epithelial surface.

The microbiome consists of both the microbiota and its host environment^74–76^. However, most microbiome studies adopt a microbe-centric approach, relying on genetic sequencing to characterize population shifts in the microbiota without assessing bacterial localization. Our findings emphasize the need to investigate host contributions to dysbiosis, particularly the role of epithelial morphology. The *Cdhr2^Δepi^* mouse model offers a platform to study bacterial communities at the epithelial surface, providing insights into environmental contributions to dysbiosis.

SFB are the most prominently visualized bacteria in the terminal ileum of *Cdhr2^Δepi^* mice. These apathogenic, spore-forming bacteria adhere to the epithelial surface via dense F-actin “holdfast points” stimulating mucosal immunity^52, 77^. Typically, SFB colonize the terminal ileum shortly after weaning and decline to low levels over several weeks^41, 42^. *Cdhr2^fl/fl^* mice showed minimal SFB presence weeks after weaning, while *Cdhr2^Δepi^* mice exhibited persistently elevated levels of SFB colonization. Disruption of the tightly packed brush border microvilli creates openings that allow bacteria like SFB to attach directly to the epithelial surface.

We observed initial SFB segments consistently co-localizing with EPS8 and Arp2/3. Arp2/3 is absent in brush border microvilli, which contain parallel, bundled F-actin, but its presence at SFB attachment sites indicates apical cytoskeletal remodeling. The pathogen, *Listeria monocytogenes*, uses Arp2/3 as an initial step to nucleate actin filaments, then augments filament elongation with the actin bunding protein Fascin^78^. Espin, consistently colocalized with SFB holdfast sites, could fulfill this role in SFB attachments. EPS8, critical for microvilli stability^16^ and actin dynamics (bundling, capping, and nucleation^16, 40, 79^), may facilitate SFB attachment by stimulating microvilli elongation and branching^80^. This is supported by EPS8’s role in nascent microvilli formation^16^ and its localization to attachment sites. Alternatively, SFB may recruit EPS8 from adjacent microvilli, destabilizing microvilli and creating larger gaps in the brush border. Data from tunneling nanotubes and spermiogenesis implicate EPS8 and Arp2/3 competitively regulate the balance between linear and branched actin networks^81–83^. Bacterial pathogens utilize bacterial effector proteins to inhibit, displace, or competitively recruit EPS8 to modify the actin cytoskeleton^11, 84, 85^. Thus EPS8 at early SFB attachments has multiple potential mechanisms to modify brush border actin structure.

SFB preferentially attach near cell-cell junctions^42^, incorporating the tight junction plaque protein^86^ ZO-1 into holdfast points in continuity with junctional complexes. Recruitment of ZO-1 suggests SFB co-opt junctional machinery potentially to trigger membrane invagination or maintain stable attachment^86, 87^. IMAC linkages maintain transcellular adhesion^28^, masking cell-cell junctions from luminal bacteria. Loss of these linkages increases SFB access to peri-junctional apical membrane surfaces, enabling attachment.

SFB have been detected in organisms ranging from chickens to primates^42, 88^, including humans^89, 90^. While SFB-like bacteria have been associated with inflammatory processes in humans^91, 92^, their role in disease remains unclear^93^.

SFB serve as a representative model for studying adherent bacteria and host defense gaps that may be exploited by other bacteria. Moreover, insights into the lifecycle of fastidious organisms such as SFB may also inform strategies for culturing other difficult-to-grow gut bacteria, which comprise up to 70% of the human gut microbiome^94^. Similar to SFB, these bacteria frequently have reduced genomes^95^ and rely on spore formation for human to human transmision^96^.

In summary, disruption of IMAC linkages between microvilli tips results in the loss of normal separation between luminal bacteria and the epithelial surface in the terminal ileum. This loss facilitates colonization by both actin-dependent adherent and mucosal-associated bacteria, representing dysbiosis via altered localization rather than microbiota composition. The Cdhr2Δepi mouse model provides a valuable tool for studying commensal bacteria at the epithelial surface in vivo, contributing to a better understanding of host-microbe interactions and dysbiosis.

## STAR Methods

## KEY RESOURCES TABLE

| REAGENT or RESOURCE | SOURCE | IDENTIFIER |
| --- | --- | --- |
| **Antibodies** |  |  |
| Anti-Arp3 (ACTR3) (mouse) [1:100] | Santa Cruz | Cat# sc-48344 |
| Anti-CD3 (rat), anti-mouse, PerCP/Cyanine5.5 | BioLegend | Cat#100218 |
| Anti-CD4 (rat), anti-mouse, Brilliant Violet 785 | BioLegend | Cat# 100453 |
| Anti-CD11b (rat), anti-mouse/human, PE/Cyanine5 | BioLegend | Cat# 101210 |
| Anti-CD45 (rat), anti-mouse, Alexa Fluor 700 | BioLegend | Cat# 103128 |
| Anti-CD45R/B220 (rat), anti-mouse/human, PerCP | BioLegend | Cat# 103234 |
| Anti-CD8a (rat), anti-mouse, PE/Cyanine7 | BioLegend | Cat# 100722 |
| Anti-EPS8 (mouse) [1:200 WM; 1:400 tissue sections] | BD Transduction Laboratories | Cat# 610144 |
| Anti-ESPN (rabbit) [1:500] | Sigma-Aldrich | Cat# HPA028674 |
| Anti-IL-17A (rat), anti-mouse, APC | BioLegend | Cat# 506916 |
| Anti-INF-γ (rat), anti-mouse, Brilliant Violet 421 | BioLegend | Cat# 505830 |
| Anti-IRTKS (BAIAP2L1) (rabbit) [1:200] | Invitrogen | Cat# PA5-54365 |
| Anti-Ly-6G (rat) anti-mouse, Brilliant Violet 510 | BioLegend | Cat# 127633 |
| Anti-TCR $\gamma/\delta$ (hamster), anti-mouse, APC/Cyanine7 | BioLegend | Cat# 118144 |
| Anti-Villin (rabbit) [1:50] | Santa Cruz | Cat# sc-28283 |
| Goat anti-mouse Alexa Fluor 488 F(ab')<sub>2</sub> fragment [1:1000] | ThermoFisher | Cat# A-11017, RRID: AB_2534084 |
| Goat anti-rabbit Alexa Fluor 568 F(ab')<sub>2</sub> fragment [1:1000] | ThermoFisher | Cat# A-21069, RRID: AB_253573 |
| **Chemicals, peptides, and recombinant proteins** |  |  |
| Alexa Fluor 647 Phalloidin [1:200] | ThermoFisher/Invitrogen | Cat# A22287 |
| DAPI (4',6-diamidino-2-phenylindole) [1:200 WM] | ThermoFisher | Cat# 62248 |
| DMSO | Sigma | Cat# D8418 |
| Ethyl Alcohol, Anhydrous (200 proof) | Decon Laboratories, Inc. | Cat# 64-17-5 |
| Ethyl Alcohol, Anhydrous (200 proof) | Electron Microscopy Sciences | Cat# 15055 |
| Glutaraldehyde, 25% | Electron Microscopy Sciences | Cat# 18220 |
| HEPES Buffer | Sigma | Cat# P1851 |
| Histo-Clear | National Diagnostics | Cat# HS-200 |
| Hoechst 33342 [1:200 WM] | ThermoFisher | Cat# 62249 |
| LIVE/DEAD (L/D) Fixable Violet Dead Cell Stain Kit | Invitrogen | Cat# L34955 |
| Osmium Tetroxide | Electron Microscopy Sciences | Cat# 19112 |
| Paraformaldehyde, 16% | Electron Microscopy Sciences | Cat# 15710 |
| ProLong Gold Antifade Reagent | Invitrogen | Cat#P36930 |
| Sucrose | Fisher Chemical | Cat# S5-3 |
| Tannic Acid | Electron Microscopy Sciences | Cat# 21700 |
| Triton X-100 | Thermo Scientific | Cat# A16046.AP |
| Uranyl Acetate | Electron Microscopy Sciences | Cat# 22400 |
| **Experimental models: Organisms/strains** |  |  |
| CDHR2<sup>fl/fl</sup> mouse (CDHR2<sup>fl/fl</sup>, villin-Cre<sup>-/-</sup>) | Tyska Laboratory, Vanderbilt University | Pinette et al., 2019 |
| CDHR2<sup>Δepi</sup> mouse (CDHR2<sup>fl/fl</sup>, villin-Cre<sup>+/-</sup>) | Tyska Laboratory, Vanderbilt University | Pinette et al., 2019 |
| CDHR5 WT and KO mice | Gross Laboratory, INSERM UMR1260 | Beck et al., 2021 |

| Other |  |  |
| --- | --- | --- |
| Precision Cover Glasses, #1.5H Thickness, 012mm | ThermoFisher / THORLABS | Cat# NC1692862 / #CG15NH |
| Superfrost slides | ThermoFisher / | Cat # 12-550-15 |
| **Software and Algorithms** |  |  |
| FIJI | https://fiji.sc | N/A |
| FIJI Plugin, Z-stack Depth Color Code | https://github.com/UU-cellbiology/ZstackDepthColorCode | N/A |
| NIS AR Elements | Nikon | N/A |
| Imaris | Oxford Instruments | N/A |
| **Prism 10** | GraphPad | N/A |
| **BioRender** | Biorender | N/A |

### Mice

Animal experiments were carried out in accordance with American Veterinary Medical Association, Vanderbilt University Medical Center IACUC, and VA Medical Center Animal Care Guidelines, ACORP Animal Protocol Number V1900133. As previously described in Pinette et al.^30^, Cdhr2^tm1c/tm1c^ mice were crossed with villin-Cre^+/-^;Cdhr2^tm1c/tm1c^. The genotype in villin-expressing tissues becomes Cdhr2^tm1d/tm1d^ which is non-expressing and referred to as Cdhr2^Δepi^. Mice that do not express Cre retain the conditional allele, Cdhr2^tm1c/tm1c,^ express CDHR2, and are referred to as Cdhr2^fl/fl^. Mating pairs comprised female *Cdhr2^Δepi^*and male *Cdhr2^fl/fl^* mice (Figure S1B). To generate consistent microbial exposure from birth through weaning, paired sets of littermates were initially co-housed to establish consistent maternal microbial exposure, then separated by genotype at weaning (Supplemental Figure 1B). Mice were sacrificed per institutional guidelines at 5 to 8 weeks of age and ileum tissue was harvested. CDHR5 WT and KO mouse tissue was provided by Isabelle Gross, INSERM UMR_S1113, Université de Strasbourg, FMTS, Strasbourg, France.

### Tissue Preparation

The distal most 3 cm of the mouse small intestine was dissected and flushed with 1.5 ml of ice-cold PBS. The unfixed, “washed” tissue was sub dissected and processed for multiple uses. After unfixed tissue was collected and snap frozen, the remaining tissue was immediately placed in fixative and further sub dissection completed in fixative as outlined below.

### Microbiome Analysis

Biospecimens were collected at weaning (stool) and sack (stool and washed ileum tissue ∼5 mm x 1 cm), snap frozen using liquid nitrogen or dry ice, then stored at −80°C until further processing. Biospecimens were processed according to the PureLink Microbiome DNA Purification Kit (A29790) protocol from ThermoFisher. Purified DNA samples were submitted to the VANTAGE genomics core at Vanderbilt University for shotgun metagenomic sequencing. Quality Control analysis was performed on the DNA samples using a DNA Qubit assay.

The samples were normalized, and libraries were prepared using the Twist Biosciences kit (P/N: 104207) following a miniaturized version of the manufacturers protocol. The library quality was assessed using the Agilent Bioanalyzer and quantified using a qPCR-based method with the KAPA Library Quantification Kit (P/N: KK4873) and the QuantStudio 12K instrument.

Prepared libraries were pooled in equimolar ratios, and the resulting pool was subjected to cluster generation using the NovaSeq 6000 System, following the manufacturer’s protocols. 150 bp paired-end sequencing was performed on the NovaSeq 6000 platform targeting 10M reads per sample. Raw sequencing data (FASTQ files) obtained from the NovaSeq 6000 were subjected to quality control analysis, including read quality assessment. Real Time Analysis (RTA) Software and NovaSeq Control Software (NCS) (1.8.0; Illumina) were used for base calling. MultiQC (v1.7; Illumina) was used for data quality assessments.

Metagenomic analysis was conducted using the Illumina DRAGEN Metagenomics Pipeline (v3.5.12). The pipeline utilized the Kraken2 algorithm to carry taxonomic classification of reads, in conjunction with a MiniKraken2 (March 2020) taxonomic database. The pipeline incorporated the DRAGEN Map/Align and de-hosting function, enabling the removal of host genome sequences and the alignment of the sequence with the murine reference genome (mm10). A minimum alignment score of 50 was applied for de-hosting control. Taxonomic classification was executed by assigning taxonomic labels to DNA sequences based on k-mers within the query sequence against the reference database. The database maps k-mers to the lowest common ancestor (LCA) among all genomes. The taxa associated with k-mers, along with their ancestors, collectively formed the taxonomy tree for classification, while sequences lacking k-mers remained unclassified. Organism detection thresholds were set between 1 and 200. Krona plots, quality control (QC) metrics, and organism detection reports were generated.

The data was returned as Kraken files which were then uploaded to Galaxy^97^ for further processing. The Bracken Abundance^98^ was processed at the phylum level with a threshold of 5. The report was filtered to include only bacteria and used to create a relative abundance graph in Prism (Figure 3H). The filtered phylum level Kraken report was loaded back into Galaxy. Krakentools^99^ was used to calculate the alpha diversity (Simpson’s Diversity Index and Berger Parker Dominance Index) and then graphed using Prism (Figure 3I). For the species analysis, the Bracken Abundance was processed at the species level with a threshold of 10. The report was filtered to include only include bacteria and species detected >2% were included in Figure 3J.

### Whole Mount (WM) Tissue Immunofluorescence

The most terminal ∼1 cm of the small intestine was excised (Cdhr2^fl/fl^ and Cdhr2^Δepi^ mice), flushed with ice cold PBS, and immediately placed in 2% PFA/PBS. Tissue was sub-dissected in fixative into small ∼2 mm^3^ pieces which were transferred to fresh 2% PFA/PBS and fixed for 1.5 h. Starting with fixation, samples were gently rocked for all incubation steps. Fixed tissue was washed three times with PBS, then permeabilized 30 m in 0.2% Triton/PBST (0.1% Tween 20 in PBS). Tissue was washed three times in PBST, and blocked in 5% normal goat serum (NGS)/PBST overnight at 4°C. Following block, tissue was washed three times with PBST and stained. Primary antibodies (listed below) were diluted in 1% NGS/PBS and incubated overnight at 4°C followed by three PBST washes. Secondary antibodies (listed below) were diluted in 1% NGS/PBS and incubated 4 hours at RT in the dark, followed by three PBS washes. DAPI, Hoechst, and phalloidin were each diluted in 1% NGS or PBS and incubated 2 h<u>ours</u> at RT in the dark. Tissue was washed four times in PBS then mounted onto glass slides in ProLong Gold using acid-washed #1.5H precision coverslips (Thorlabs #CG15CH2) with the villus surface facing the glass coverslip. The following antibodies and dilutions were used for whole mount staining: anti-EPS8 (mouse BD Transduction Laboratories #610144), 1:400; anti-Villin (rabbit, Santa Cruz #sc-28283), 1:50; anti-ESPN (rabbit, Sigma-Aldrich #HPA028674), 1:500; anti-Arp3 (ACTR3) (mouse, Santa Cruz #sc-48344), 1:100; Goat anti-mouse Alexa Fluor 488 F(ab’)2 fragment (Invitrogen #A-11017), 1:1000; Goat anti-rabbit Alexa Fluor 568 F(ab’)2 fragment (Invitrogen #A-21069), 1:1000; Phalloidin 647 (Invitrogen #A22287), 1:200; and DAPI (ThermoFisher #62248), 1:200; and Hoechst 33342 (ThermoFisher #62249), 1:200. Secondary antibodies (not including phalloidin, DAPI, or Hoechst) were centrifuged at 4°C for 10 m at a speed of 21,000 x g prior to use.

### Frozen Section Tissue Preparation

Tissue was placed in 2% PFA and further sub dissected in fixative into pieces ∼2mm^2^, then fixed for an additional 30 minutes at RT, total fixation time ∼60 minutes. After fixation, the tissue samples were washed three times with PBS. Samples were placed in 30% sucrose villi-side down and incubated overnight at 4°C. Then samples were then washed in optimal cutting temperature (OCT) compound by passing the tissue through 3 separate blocks of OCT. Samples were oriented will villi pointing laterally, perpendicular to the lab bench, in a fresh block of OCT and snap frozen in dry ice-cooled ethanol. Using cryostat Leica CM2050-S samples were cut into 10µm thin sections and mounted on Superfrost Plus Microscope Slides (fisherbrand) and stored at −20°C until staining.

### Frozen Section Immunofluorescence

Slides were thawed to RT and washed twice with PBS. Samples were permeabilized using 0.2% Triton X-100 in PBS for 30 minutes at RT. Coverslips were washed with PBS, blocked in 5% normal goat serum (NGS) in PBS overnight at 4°C in a humidified chamber. Following block, tissue was washed three times with PBS. Primary antibodies (listed below) were diluted in 1% NGS/PBS and incubated on the samples overnight at 4°C in a humidified chamber followed by three PBS washes. Secondary antibodies (listed below) were diluted in 1% NGS/PBS and incubated 4 h at RT in a dark humidified chamber, followed by three PBS washes. DAPI, Hoechst, and phalloidin were each diluted in 1% NGS or PBS and incubated 2 h at RT in the dark. Finally, samples were washed 4 times with PBS and mounted in ProLong Gold using acid-washed #1.5H precision coverslips (Thorlabs #CG15CH2). Samples were left in a dark chamber at RT overnight to cure. The following antibodies/stains and dilutions were used: Anti-EPS8 (mouse, BD Transduction Laboratories Cat# 610144), 1:200; Anti-E-cadherin (mouse, BD Transduction Laboratories Cat# 610182), 1:100; Anti-ZO-1 (rabbit, Invitrogen #61-7300), 1:50; goat anti-mouse Alexa Fluor 488 F(ab’)2 fragment (Invitrogen Cat# A-11017), 1:1000; goat anti-rabbit Alexa Fluor 568 IgG (H+L) (Invitrogen Cat# A-11011), 1:500; Alexa Fluor 647 Phalloidin (Invitrogen Cat# A22287), 1:200. Secondary antibodies were centrifuged at 4°C for 10 m at a speed of 21,000 x g prior to use. Samples mounted in ProLong Gold and allowed to cure before imaging.

### Paraffin Section Tissue Preparation

After washing with ice cold PBS, mouse ileum was placed in 2% PFA and further sub dissected into pieces ∼2 mm^2^. After a total of 2 hours in fixative, samples were placed in a tissue cassette, transferred to 70% EtOH and submitted to the Vanderbilt University Translational Pathology Shared Resource to be embedded in paraffin wax. Using a Microm HM-325 microtome samples were cut into 7-15 µm sections and mounted on glass slides and stored at RT until staining.

### Paraffin Section Immunofluorescence

Paraffin slides were de-paraffinized by placing on a 60°C slide warmer followed by two Histoclear washes. Slides were rehydrated through an ethanol gradient (EtOH 100% x2, 95% x2, 70%), then 20 mM Tris-HCl, pH8. After rehydration, samples were processed as above for Frozen Section Immunofluorescence.

### Fluorescent *in situ* Hybridization (FISH)

Samples were hybridized with FISH probes after permeabilization/blocking but before primary and secondary antibody staining as follows. Samples were incubated with FISH probe SFB1008-CY3 (5’-Cy3-GCGAGCTTCCCTCATTACAAGG-3’, Sigma Genosys Custom Oligo)^43, 100^ at 2 µM in Hybridization Buffer (20 mM Tris-HCl pH 8.0, 0.9 M NaCl, 0.01% SDS) in a dark, humidified, hybridization chamber for 2 hours at 46°C. Samples were then washed in FISH Wash Buffer (20 mM Tris-HCl pH 8.0, 225 mM NaCl, 5 mM EDTA). Following wash step, samples were incubated with antibody or other stains as out lined above. FISH probes were centrifuged at 4°C for 10 m at a speed of 21,000 x g prior to use.

### Electron Microscopy

Steps are completed at room temperature (RT) unless indicated otherwise. Following wash with PBS (phosphate buffered saline), tissue is placed in 4% paraformaldehyde (PFA)/PBS and sub-dissected into small pieces ∼5 mm or less using a stereo dissecting microscope, then transferred to a large volume (5-10 mL) of EM fixative (2.5% glutaraldehyde, 4% PFA in SEM Buffer to complete fixation 1 h. SEM Buffer consisted of either PBS or 0.1M HEPES, pH 7.3 with subsequent washes in the corresponding Buffer (without Ca). Samples were washed 3x in SEM Buffer, then incubated in 1% tannic acid/SEM Buffer for 1 h. Tissue was washed 3x with ddH_2_O followed by 1 h incubation with 1% osmium tetroxide/ddH_2_O. Tissue was washed three times with ddH_2_O, then incubated 30 m with 1% uranyl acetate/ddH_2_O. Samples were washed three times in ddH_2_O, then dehydrated in a graded ethanol series and dried using critical point drying (*Cdhr2^fl/fl^* and *Cdhr2^Δepi^*mouse tissue samples) or chemically with hexamethyldisilazane (*Cdhr5* WT and *Cdhr5* KO mouse tissue samples). Samples were mounted on aluminum stubs and sputter coated with platinum (*Cdhr2^fl/fl^* and *Cdhr2^Δepi^* samples) or gold/palladium (*Cdhr5* WT and *Cdhr5* KO tissue samples) using a Cressington 108 sputter coater. Imaging was performed using a Quanta 250 Environmental SEM operating in high-vacuum mode with an accelerating voltage of 5-10 kV (Figure 1B-E, 5A) or Zeiss Crossbeam 550 operated at 2 kV using the lower secondary electron detector (Figure 4G, 5B).

### Fluorescence Microscopy (Whole Mount and Tissue Sections)

Laser scanning confocal microscopy of whole mount samples was performed on a Nikon FN1 upright light microscope equipped with an A1 R HD25 laser-scanning head, four excitation LASERs (405, 488, 561, and 647 nm), and a 60x/1.4 NA objective. All images used for quantitative comparisons were prepared with equal treatment, acquired with identical parameters (e.g. pinhole diameter and detector gain), and processed in an identical manner.

Spinning disk confocal microscopy of tissue sections was performed using a Nikon Ti2 inverted light microscope with a Yokogawa CSU-W1 spinning disk head with SoRA, a Hamamatsu Fusion BT sCMOS camera, and four excitation LASERs (405, 488, 568, and 647 nm). A 60x/1.49 NA TIRF oil immersion or 100x/1.45 NA oil immersion objective was used. To visualize protein localization at individual bacteria attachment sites, we performed confocal imaging with oversampling followed by deconvolution.

### Image Processing

Images were deconvolved using Nikon Elements. Image stacks were viewed as single confocal planes, 3D renderings (Nikon Elements or Imaris), or as a maximum intensity projection (maxIP) generated in ImageJ/FIJI, Nikon Elements, or Imaris. Images were contrast-enhanced and cropped using a combination of ImageJ/FIJI software from the National Institutes of Health (NIH), Nikon Elements, and Imaris. Post-processing to segment DAPI images into cellular nuclei and bacterial DNA was completed using (a) Labkit pixel classification^101^ with ImageJ/FIJI, (b) Labkit pixel classification with ImageJ/FIJI plus Imaris or (c) Imaris built-in machine learning segmentation. Once segmented, the cell nuclear DNA and bacterial DNA were viewed in separate channels and differentially contrasted to facilitate visualization of the bacterial DNA signal. Imaris was used to generate 3D surfaces representing brush border actin and surface bacteria (Figure S3). Dual-depth-coded images were generated as separate images in ImageJ/FIJI using the “Z-stack Depth Color Code” plugin then combined into a single image as a MaxIP (Figure 4).

### RNA Isolation and Real Time qRT-PCR

RNA was isolated from washed ileum tissue from paired sets of *Cdhr2^fl/f^*and *Cdhr2^Δepi^* littermates at 5 to 8 weeks of age using TRIzol (Invitrogen) as described by the manufacturer followed by an RNA clean up protocol using the Qiagen RNAeasy kit. RNA was transcribed to cDNA using Thermo Fisher Scientific HighCapacity cDNA Reverse Transcription Kit (ThermoFisher Scientific, Waltham, MA, USA) per the manufacturer’s specifications and then Taqman based real time rtPCR was performed using TaqMan Fast Advanced Master Mix (ThermoFisher Scientific, Waltham, MA, USA) and Taqman gene expression assays on the Applied Biosystems QuantStudio 6 Flex instrument. Relative Units were quantified using the relative gene expression. In short, all samples were calibrated to a pooled control sample prepared from control mice. Relative units were calculated using the 2^^-ΔΔCt^ method, where the ΔCt is calculated as the difference in the cycle threshold of the gene of interest from the endogenous control gene (*Gapdh*) and then the ΔΔCt is the difference between the ΔCt of the sample compared to the control sample. Primer and probe sets were purchased as Taqman Gene Expression Assays from Applied Biosystems [mouse primer sets: *Cxcl1* Mm04207460_m1, *Il22* Mm01226722_g1, *Il6* Mm00446190_m1, *Il17a* Mm00439618_m1, *Reg3g* Mm00441127_m1 and *Gapdh* Mm99999915_g1].

#### Method for Flow Cytometry

Single-cell suspensions were prepared from mouse spleens and lymph nodes by mechanical dissociation through 40 µm strainers into T cell media, followed by RBC lysis (1 mL RBC lysis buffer, 1 min, quenched with 20 mL PBS + EDTA) and centrifugation (350 × g, 7 min, 4 °C). Cells were stained with Fixable Violet Dead Cell Stain (Invitrogen, #L34955) diluted 1:1000 in PBS for 30 min at 4 °C in the dark, washed, and surface-stained in 100 µL MACS buffer for 30 min at 4 °C with CD45-AF700 (BioLegend #103128), CD3-PerCP-Cy5.5 (#100218), CD4-BV785 (#100453), CD8-PE-Cy7 (#100722), γδT-APC-Cy7 (#118144), B220-PerCP (#103234), Ly6G-BV510 (#127633), and CD11b-PE-Cy5 (#101210). Cells were fixed with Reagent A (Fix & Perm Kit, Thermo Fisher, Cat# GAS003) for 20 min at RT, washed, and permeabilized in Reagent B containing IL-17A-APC (#506916) and IFNγ-BV421 (#505830) for 30 min at RT in the dark. Fluorescence minus one (FMO) controls contained all antibodies except the omitted parameter, and single-color compensation controls were prepared with antibody capture beads. Data were acquired on a Cytek Aurora flow cytometer and analyzed using standard gating strategies.

## QUANTIFICATION AND STATISTICAL ANALYSIS

### Ratio of bacterial F-actin attachments to cell nuclei quantification

Using ImageJ/FIJI, the standard depth in Z analyzed was 50-150µm to account for the variable angles and positions of villi in a sample. Villi positioned at an angle were analyzed in increments down the Z plane to keep the consistent depth of 50-150µm. DAPI signal was used to quantify the number of nuclei within the selected depth. F-actin attachments perpendicular to the Z plane were identified by visualization of a circular ring that constricted to a point as the Z plane moved from surface to within the villus. Horizontal F-actin attachments were identified by visualization of triangular inclusions along the villus edge when parallel to the Z plane. To accurately visualize F-actin attachments throughout the selected depth the brightness and contrast of the image were adjusted for each plane. The sums of both the nuclei and F-actin attachments were used to calculate the ratio. The same approach was taken quantify the F-actin attachments positive for EPS8.

### Bacteria DNA Pattern Quantification

To quantify the bacteria DNA patterns observed in sectioned tissue, the previously quantified sums of bacteria documented in Figure 2G/H were used as the total bacteria in a sample. Then images were evaluated across multiple z-steps to determine if bacteria were single, filamentous or indeterminate (most commonly due to high concentrations of bacteria). Quantification was blinded for the source of tissue (*Cdhr2^fl/fl^*or *Cdhr2^Δepi^)*.

### Distance from Cell Junction Quantification

Confocal images were opened in ImageJ/FIJI and the measuring tool was used to record the distance from the center of the bacteria to the closest visible cell junction.

### Analysis of Signal Intensity at Bacterial Attachment Sites

To measure signal intensities at bacterial attachment sites, 1-pixel wide line scans were drawn either transverse parallel to the cell surface or longitudinal from intracellular through the attachment site to extracellular. All intensity values were normalized (range set from 0 to 1). The intensity was normalized for each channel and fit to a Gaussian curve or a sum of 2 Lorentzian curves using PRISM v. 10.6.1 (GraphPad).

### Statistical Analysis

Statistical significance was performed using the Mann-Whitney test (Figure 2G-H, Figure 5C), Friedman test (Figure 3I), unpaired T test (Supplemental Figure 1C), and paired Wilcoxon test (Supplemental Figure 1F-G). All statistical analysis was computed in PRISM v. 10.6.1 (GraphPad).

## ACKNOWLEDGEMENTS

The authors would like to thank members of the VUMC EBC for constructive feedback. We acknowledge the Translational Pathology Shared Resource (NCI/NIH Cancer Center Support Grant P30CA068485), Microscopy was performed in part through the Vanderbilt Cell Imaging Shared Resource (supported by NIH grants CA68485, DK20593, DK58404, and EY08126). SEM images were collected on a FEI Quanta 250 Scanning electron microscope funded by S10 RR026373 or Zeiss 550 Crossbeam FIB-SEM funded by S10 OD028704. This work was supported by the Department of Veterans Affairs Career Development Award 1IK2BX004885 (L.M.M.).

---

**Supplemental Figure S1.**
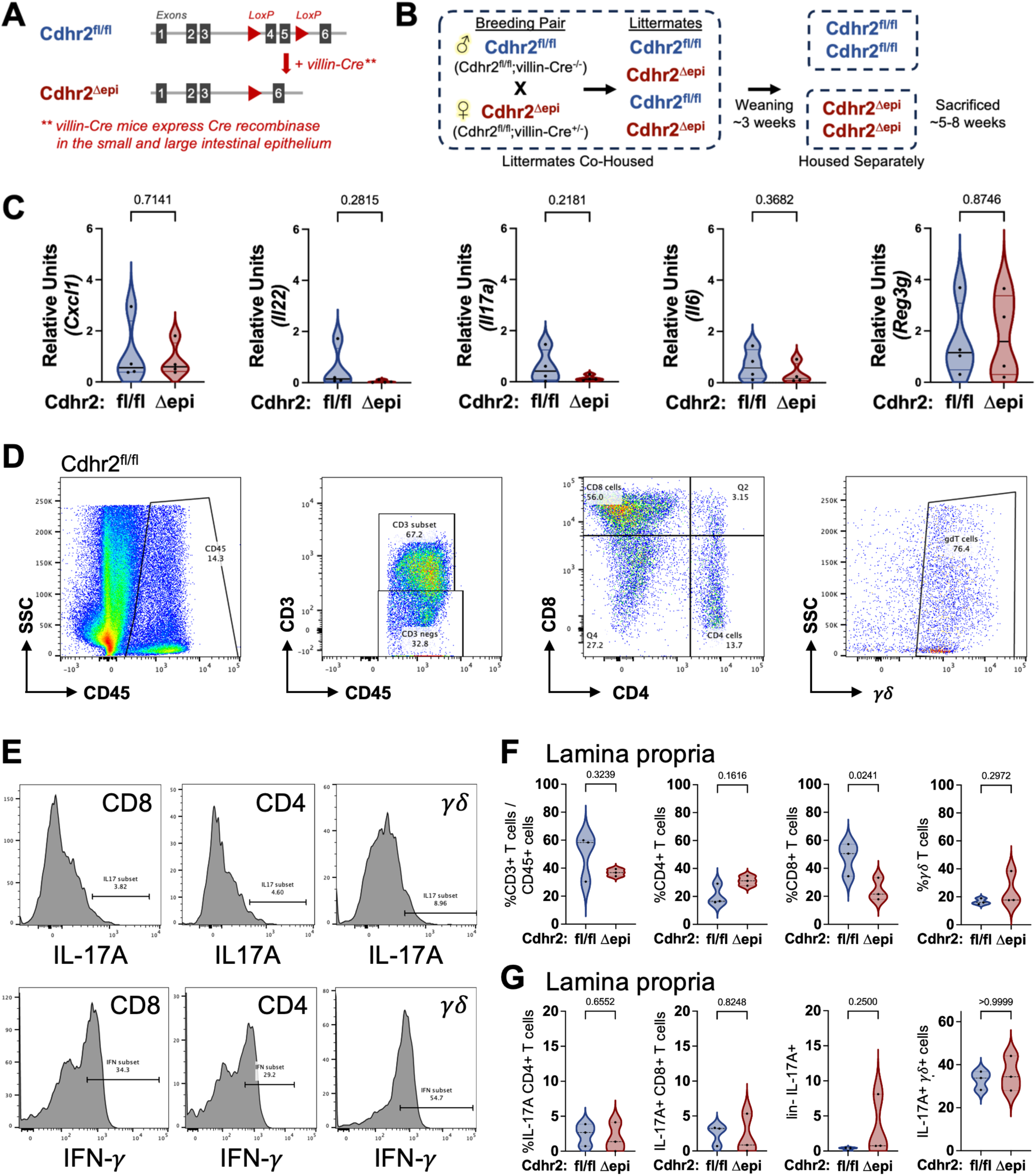
Mice with a disrupted IMAC do not have increased inflammation. (A) Simplified diagram of the strategy used to generate *Cdhr2*^Δepi^ mice from *Cdhr2^fl/fl^* in combination with tissue-specific expression of Cre; previously described in Pinette et al. 2019^30^. (B) Diagram of breeding strategy. Using a background of *Cdhr2^fl/fl^* mice, introduction of a single copy of villin-Cre (*villin-Cre^+/-^*) in the mating pair results in a mixed litter of *Cdhr2^fl/fl^* and *Cdhr2^Δepi^* mice. (C) The expression of *Cxcl1, Il22, Il17A, Il6* and *Reg3g* in RNA extracted from the terminal ileum at 6-8 weeks of age was assessed by qRT-PCR. Relative units are calculated as described in the Materials and Methods, *Gapdh* was used as endogenous control. Statistical analysis performed using unpaired t-tests. Error bars represent mean ± SEM. (D-G) Flow cytometry analysis of T cells within the lamina propria in knockout C*dhr2^Δepi^* and control *Cdhr2^fl/fl^*mice. (D) Two-dimensional flow cytometry plots show the gating strategy of CD45^+^ immune cells in gastrointestinal tissue to identify CD3^+^ T cells within CD45^+^ immune cells, CD4^+^ and CD8^+^ T cells and γδ T cells. (E) Histograms show gating strategy used to identify IL-17A and IFN-γ expressing cells in CD8^+^ T cells, CD4^+^ T cells, and γδ T cells. (F) Violin plots show proportions of T cell populations in the lamina propria of matched *Cdhr2^fl/fl^*and *Cdhr2*^Δepi^ mice (CD3^+^ T cells over total CD45^+^ T cells) and (CD4^+^ T cells, CD8^+^ T cells and γδ T cells over total CD3^+^ T cells). (G) Violin plots show proportions of IL-17A expressing T cells (CD4, CD8), lineage negative and γδ T cells, in the small intestine lamina propria of matched *Cdhr2^fl/fl^* and *Cdhr2^Δepi^*mice, n = 3. Statistical analysis performed using paired Wilcoxon test.

**Supplemental Figure S2.**
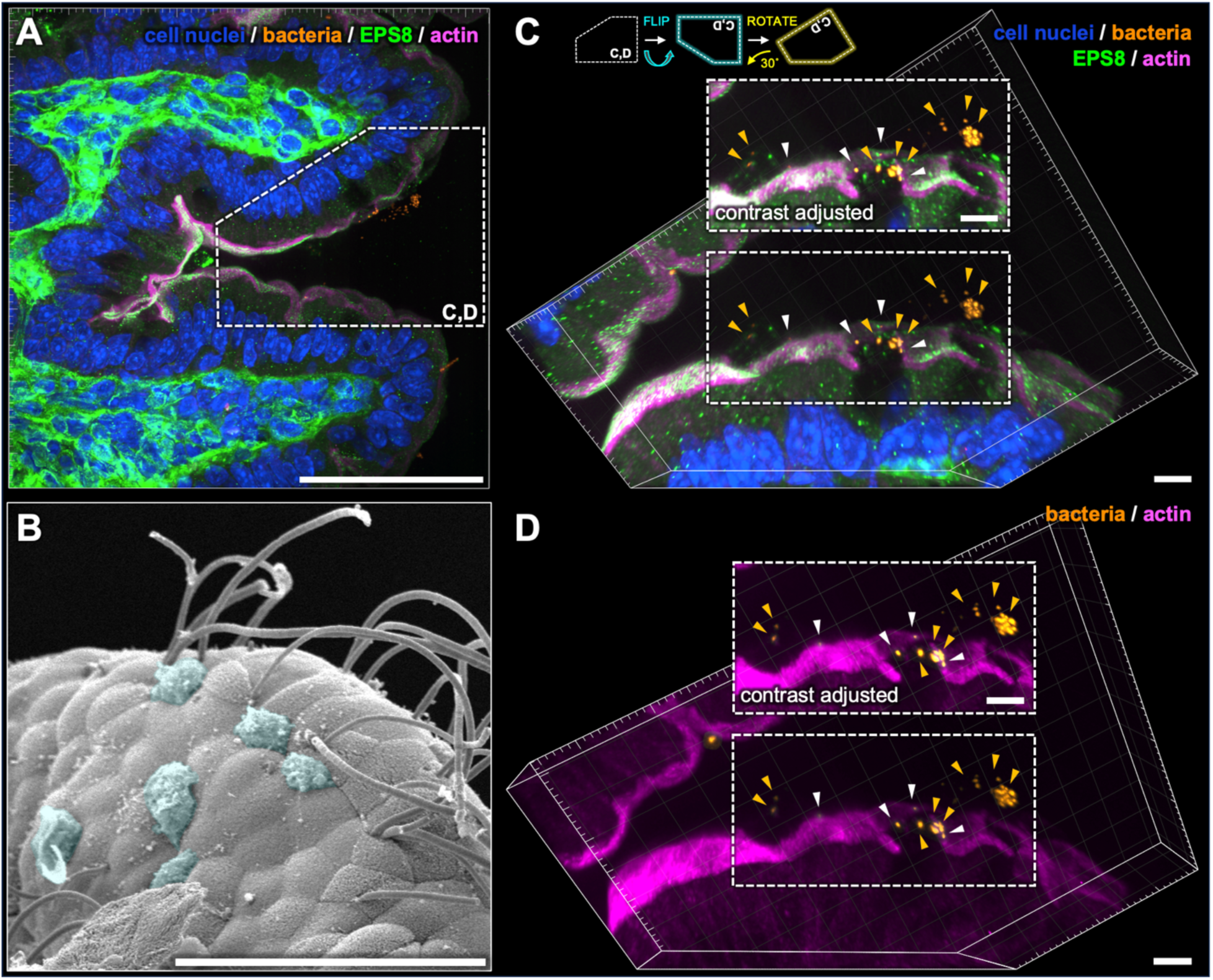
Tissue-associated bacteria form actin-independent surface attachments—goblet cell and mucus binding. (A) Deconvolved confocal 3D image of *Cdhr2^Δepi^* ileum viewed as a maxIP (depth 11.25 µm) stained for DNA (DAPI), EPS8, and actin. DAPI segmented using an AI algorithm to isolate and differentially contrast bacteria (orange) relative to cell nuclei (blue). Boxed area enlarged and rotated in C and D. (B) SEM of *Cdhr2^Δepi^*ileum. Retained mucus highlighted by transparent cyan overlay. (C) Boxed area from (A) with cell nuclei, bacteria, EPS8, and actin; cropped, enlarged, contrast enhanced, and positioned (flipped vertically, rotated ∼30° counterclockwise, tilted) to optimize visualization of bacteria relative to the cell surface. Arrowheads highlight bacteria in contact with goblet cells (white) or the adjacent lumen (orange). Boxed area in shown above with augmented brightness/contrast to visualize dim signals. (D) Boxed area from (A) showing only the bacteria and actin channels; cropped, enlarged, contrast enhanced, and positioned (flipped vertically, rotated ∼30° counterclockwise, tilted) to optimize visualization of bacteria. Arrowheads highlight bacteria in contact with goblet cells (white) or in the adjacent lumen (orange). Boxed area in shown above with augmented brightness/contrast to visualize dim signals. Scale bars: 50 µm (A, B), 5 µm (C, D).

**Supplemental Figure S3.**
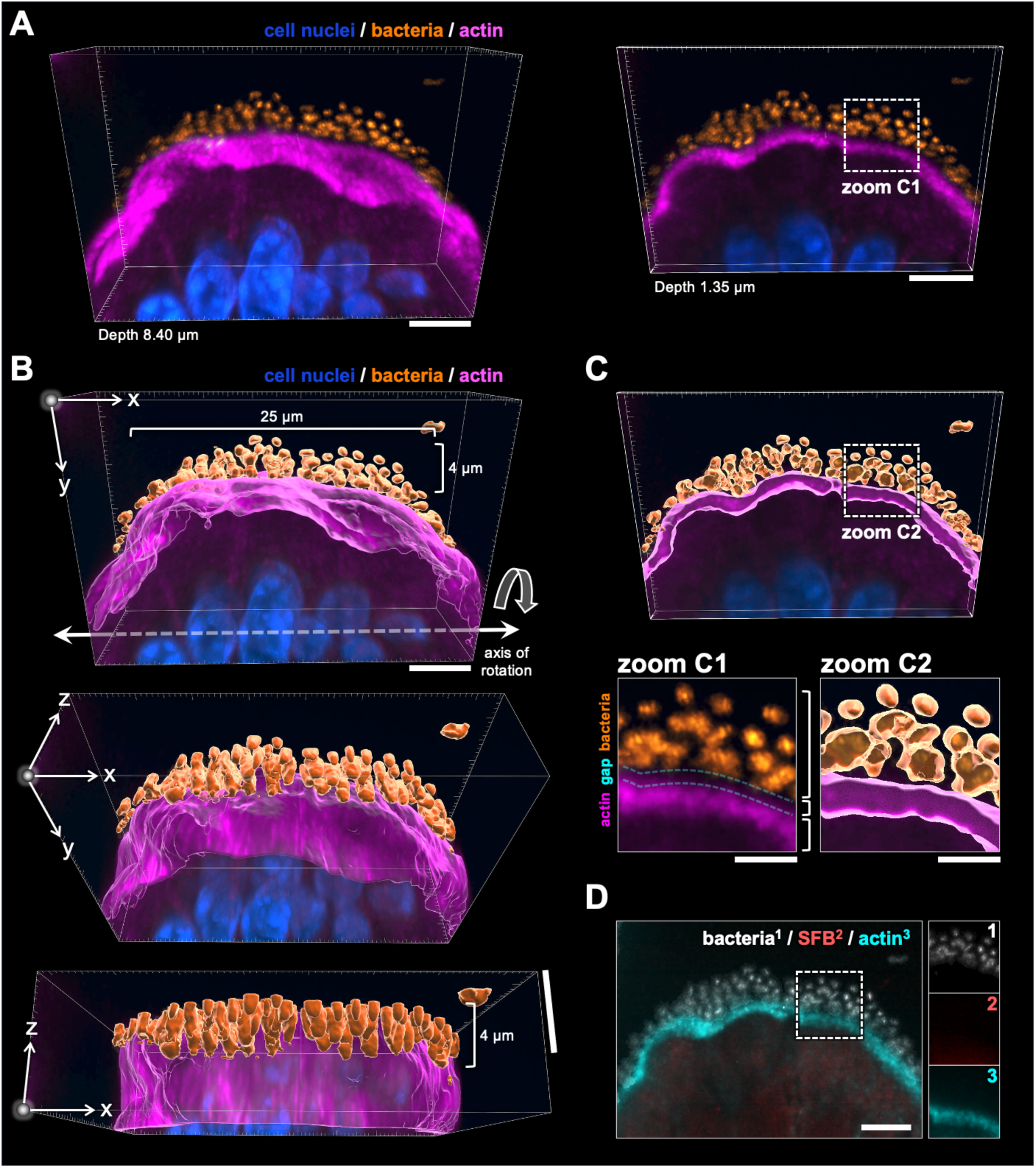
Mucosal bacteria form layered three-dimensional structures that are actin-independent. (A) Villus tip from Cdhr2^Δepi^ ileum (Figure 3J, zoom 1) viewed as a MaxIP (depth 8.40 µm, left; depth 1.35 µm, right) and rendered in 3D. Tissue stained with DAPI (DNA) and for actin. DAPI segmented using an AI algorithm to isolate and differentially contrast bacterial DNA (orange) relative to cell nuclei (blue). Boxed area enlarged in zoom C1. (B) MaxIP (depth 8.40 µm) from (A) processed using Imaris to create a 3D surface from the fluorescent actin and bacteria DNA signals. The surface is overlaid on the fluorescence image, then sequentially rotated around the x-axis to highlight the 3D composition of the bacteria overlying the F-actin surface. (C) MaxIP (depth 1.35 µm) from viewed as a surface to highlight the relationship between the bacteria and surface actin. Boxed area enlarged and contrast enhanced in zoom C2. Zooms C1-2 show enlarged images of the original fluorescence intensity (C1) and generated surface (C2) with a clear gap between the bacteria fluorescence and surface actin and continuity of the actin surface. (D) MaxIP (depth 1.35 µm) corresponding to the image volume in (A, right) with segmented DNA (bacteria), SFB FISH (SFB), and actin. Boxed area shown as separate channels (right). Scale bars: 5 µm (A-D), 2 µm (zoom C1-2).

